# PTPN1/2 inhibition induces highly functional terminal effector CD8 T cells through autocrine IL-10

**DOI:** 10.1101/2023.04.17.537264

**Authors:** Luis-Alberto Perez-Quintero, Alexandre Poirier, Zuzet Martinez-Cordoba, Chu-Han Feng, Isabelle Aubry, Cedric Carli, Samaneh Kamyabiazar, Alain Pacis, Yevgen Zolotarov, Kelly-Anne Pike, Jean-Sebastien Delisle, Michel L. Tremblay

**Author notes:** Correspondence: Dr. M. L. Tremblay, Ph.D., Department of Biochemistry, Microbiology, and Immunology, Rosalind, and Morris Goodman Cancer Centre, McGill University, 1160 Avenue des Pins, Montreal, Quebec, Canada H3A1A3.

## Abstract

Increased understanding of the modulatory pathways controlling CD8 T cell responses has led to the formulation of successful checkpoint inhibitor-based immunotherapies against cancer. However, their effectiveness is limited to a few tumor types, motivating the search for novel combinatorial strategies. PTPN1 and PTPN2 are two homologous protein tyrosine phosphatases recently proposed as potent intracellular checkpoints. Furthermore, their catalytic domain is a propitious target for small-molecule pharmacological intervention. Herein we investigated the potential effects of conditional genetic deletion of either or both phosphatases in mouse CD8 T cells, one of the main effectors in cancer immunotherapy. Our results demonstrated that hemizygous deletion of PTPN1 in a PTPN2 deficient background heightens the enhanced effector phenotype already observed in PTPN2 defective CD8 T cells. This functional gain is mediated by an autocrine IL-10 positive feedback loop. Pharmacological inhibition with a PTPN1/2 small-molecule inhibitor yielded similar results, highlighting the importance of simultaneously inhibiting both phosphatases. Our study uncovers a novel mechanism by which the downregulation of PTPN1 and PTPN2 act as a powerful tool for potentiating CD8 cytotoxic responses.

## Introduction

PTPN1 and PTPN2 are two highly homologous nonreceptor protein tyrosine phosphatases with preferential activity on targets of the JAK/STAT pathway [1]. Recent studies by us and others have underscored the potential of these phosphatases as immunotherapeutic targets [2–6]. Indeed, small molecule inhibitors targeting both phosphatases have shown strong synergy in syngeneic mouse models partially responding to anti-PD1 [7, 8] and currently are been tested in Phase I trials by a Calico-AbbVie consortium (Trial IDs NCT04777994 and NCT04417465). Phylogenetically, both phosphatases derive from a common ancestor [9, 10] and map closely in an independent branch of the PTP family [11]. This identity reaches up to 70% in the catalytic domain [12] and manifests functionally by sharing common substrates. Mechanistically, PTPN1, and PTPN2 present a pronounced substrate recognition motif involving tandem phosphotyrosines [13], hence the overlap in their targeting molecules, mainly belonging to the receptor tyrosine kinases (RTK) [14, 15], JAK/STAT [16–20], and Src [21] families. Attempts to genetically block the expression of both enzymes simultaneously highlighted a potential overlapping role, particularly in the dephosphorylation of STAT1 [22]. Embryonic lethality of PTPN1 and PTPN2 double deletion in mice, strongly supports overlapping roles since single mutant animals for any of both enzymes, reach birth at mendelian rates [23, 24]. However, their differential expression, cellular localization, and substrate preference are indicative that the redundancy is partial [1]. Pharmacologically, the similarities between the catalytic domains of PTPN1 and PTPN2 have challenged the rational design of small molecule inhibitors with selectivity for either of these enzymes [25–27]. Nevertheless, the potential benefits of restricting the activity of both phosphatases in tumor immunotherapy opens a convenient door for the use of dual-targeting small molecule inhibitors of PTPN1 and PTPN2[2, 7, 8, 28].

CD8 T cells are essential immune system components in mounting cellular immune responses [29]. Increasing their ability to function even in immunosuppressive microenvironments has been a major challenge for tumor immunotherapy. Hence, PTPN1 and PTPN2 offer a rational alternative to the current known immune checkpoint inhibitors which act by the recruitment of two alternative phosphatases, SHP-1 and SHP-2 [28]. Indeed, reports with PTPN1 and PTPN2 inhibitors establish T cells as the one of the main immunological targets of these compounds[7, 8].

Interleukin 10 (IL-10) is a pleiotropic cytokine regarded mainly for its regulatory roles on myeloid and CD4 T cells [30–32]. However, it can induce proinflammatory signals in a timely and cell dependent manner [33]. In CD8 T cells, IL-10 had shown importance in maintaining cytotoxic potential and formation of memory in vivo [30, 34], in enhancing CD8 mediated immunopathology in the liver [35] and in inducing the activation of tumor-resident CD8 T cells [36, 37]. IL-10 signals through different components of the JAK/STAT pathway, particularly STAT3 [38, 39], which is regulated by PTPN1 [40]. Indeed, STAT3 activity, either by IL-10 or IL-27 stimulation, has been shown to be required for the differentiation of highly functional TIM3+ and CX3CR1+ terminal effector CD8 T cells in infectious and tumor models [41–43].

Herein, we analyzed CD8+ cells from mice with conditionally targeted deletions of PTPN2 and PTPN1. Whereas we found an enhanced functional state in PTPN2 single deficient cells, additional hemi-deficiency of PTPN1 greatly enhanced their pro-cytotoxic abilities which was dependent on an autocrine IL-10 feedback loop. Moreover, small molecule inhibitors targeting both enzymes replicated this mechanism, contributing to the overall understanding of their pharmacological effects.

## Results

### Combined deletion of PTPN1 and PTPN2 alters CD8 T cell differentiation

To address the combined contribution of PTPN1 and PTPN2 to the regulation of activating signals in CD8+ T cells, we designed a mouse line where allelic inactivation of either one or both genes occurs in mature, post-thymic CD8 T cells. To achieve this, we breed heterozygous mice carrying “floxed” alleles of both genes, PTPN2 [44] and PTPN1 [45], with a mouse line where the CRE recombinase expression is driven by the CD8 enhancer 8I (E8i-CRE) [46]. Mice were bred to produce littermates with either WT (wt/wt), heterozygous (wt/fl) or homozygous (fl/fl) versions of either single or both genes (Figure S1A). To avoid potential artifacts arising from off-target effects of the CRE recombinase [47, 48], mice heterozygous for E8i-CRE (wt/cre+) carrying wt alleles for PTPN1 and PTPN2 were used as controls, hereafter named CRE. Mice were born at Mendelian rates, and no apparent differences were detected between the various genotypes until 6-8 weeks of age. CD8 T cells from all genotypes reached maturity and were distributed in peripheral secondary immune organs (Figure 1A). To confirm that conditional deletion of both genes was achieved, we tested the protein expression of PTPN2 and PTPN1 by immunoblotting in both, peripheral CD4 and CD8 T cells (Figure S1B). As expected, protein expression corresponded to each genotype, and deletion was not observed in CD4 T cells. Nevertheless, mice around 8-10 weeks of age in which CD8+ T cells lacked expression of both genes (ptpn1 fl/fl; ptpn2 fl/fl; wt/cre+) or dKO hereafter, showed a significant reduction in thymus size with complete involution observed around 12 weeks of age (Figure S1C). Microscopically, cortical, and medullary architecture was lost in the thymi of these mice when compared to CRE controls (Figure S1D). The thymic involution was accompanied by the onset of lesions compatible with scaly skin disease, an opportunistic *Corynebacterium bovis* infection observed in other strains of athymic mice (Figure S1E) [49]. Lesions were present only in exposed skin and were characterized by ulceration, neutrophilic inflammation, acanthosis, and hyperkeratosis/crusting. dKO mice were routinely euthanized based on humane grounds as the skin lesions were severe enough to cause dehydration, subsequent loss of weight and ultimately death. This phenotype was observed with 100% penetrance in dKO mice by 12 weeks of age, conversely, it was not observed in other genotypes studied. dKO mice presented also mild splenomegaly and spontaneous lymphadenopathies (Figure S1F). In contrast to dKO mice, other genotypes did not show differences in organ size or cellularity when compared to CRE controls (Figure S1F).

**Figure 1.**
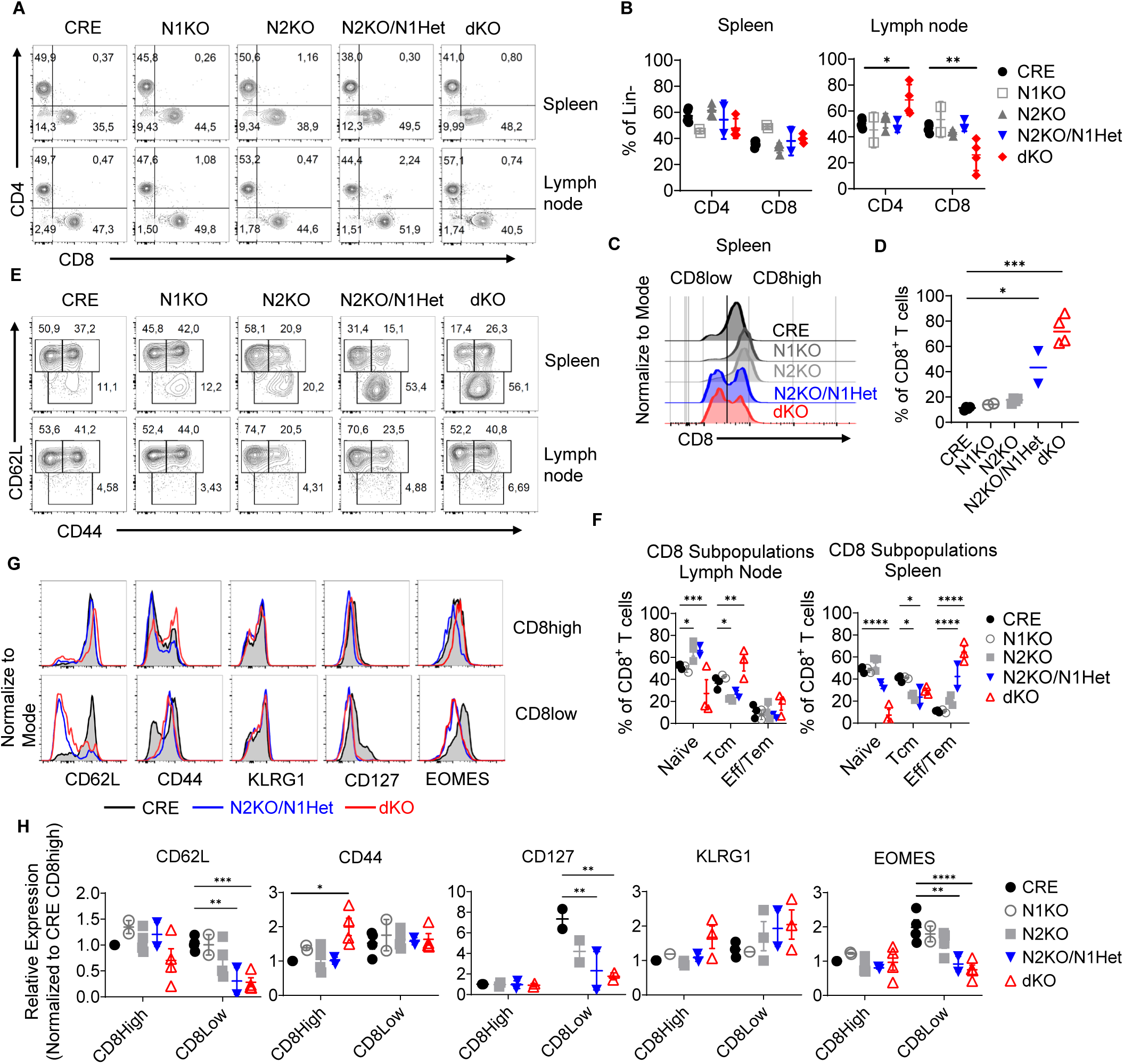
Post thymic deletion of PTPN2 and PTPN1 in CD8 T cells increases their activation and differentiation in naïve mice. Ex vivo analysis by multiparametric flow cytometry of peripheral T cell populations of naïve 6 to 8 weeks old mice with conditional deletions of *Ptpn1* and/or *Ptpn2* in mature post-thymic CD8 T cells. A) Representative CD4 vs CD8 distribution from different genotypes in spleen and inguinal lymph nodes. Events shown correspond to single, viable, and lineage-negative (Lin-) gates. B) Percentages expressed as mean and S.E.M. of data from different experiments C) Histogram representation of CD8 expression level in splenic single positive CD8 T cells and D) percentages of CD8 low populations observed within in the different genotypes. E) Plots and F) Statistical representation of CD8 T cell CD62L and CD44 expression from spleen and lymph nodes; naïve cells are defined as CD62L high and CD44 low, T central memory (Tcm) defined as CD62L high and CD44 high cells, and T effector/effector (Teff/eff) memory as CD62L low and CD44 intermediate cells. G) Histogram representation and H) Quantification of differentiation markers EOMES, KLRG1, and CD127. (CRE n=4, N1KO n=2, N2KO n=4, N2KO/N1Het n=2, dKO n=4) (Bars represent +/- S.E.M.; non-parametric Kruskal-Wallis test (D) or Mixed-effects analysis (B, F, H); P values *<0.05, **<0.005, ***<0.0005, ****<0.00005).

Next, we characterized circulating CD8 populations from the different genotypes by flow cytometric analysis. Despite CD8 T cells being present in the spleen and lymph nodes of all the genotypes analyzed, mice carrying dKO CD8 T cells showed an increase in the CD4/CD8 cell ratio in lymph nodes, however, the absolute number of CD8 T cells was not significantly affected (Figures 1A, 1B and S1F). This finding, together with the size increase of lymph nodes, suggested a change in the migratory cues of dKO circulating CD8 T cells. No other significant differences in the distribution of T cells in lymph nodes or spleen were noted in the genotypes analyzed. Nevertheless, we detected an increase in the subpopulation of splenic CD8 T cells expressing low levels of the CD8 coreceptor in both, dKO and N2KO/N1Het (Ptpn1 wt/fl; Ptpn2 fl/fl; wt/cre+) mice (Figures 1A and 1C). Accumulation of these CD8low-expressing cells was more pronounced in the dKO animals, almost doubling those seen in the N2KO/N1Het animals, comprising around 70% and 40% of all splenic CD8 T cells, respectively (Figures 1C and 1D). Cells displaying low expression of the CD8 coreceptor were also observed in CRE controls, and both single KOs, yet differences between them and CRE controls were not statistically significant (Figures 1C and 1D). To further characterize peripheral CD8 T cells in these animals, we assessed their differentiation state by co-staining for the trafficking molecule L-selectin (CD62L) and the memory marker CD44 (Figure 1E). We sorted different CD8 T cell subpopulations of the spleen and peripheral lymph nodes based on CD62L and CD44 surface molecule expression defining naïve, effector/effector memory, and central memory subpopulations (Figure 1E and 1F). While the differentiation state of CD8 T cells in lymph nodes seemed unaffected within the various genotypes, spleens showed a marked increase of effector/effector memory-like CD8 T cells in the N2KO/N1Het mice (Ptpn1 wt/fl; Ptpn2 fl/fl; wt/cre+) and dKO mice. This subset of CD8+ T cells, characterized by reduced expression of CD62L and intermediate expression of CD44, correlated with the presence of CD8low cells (Figure 1D). To determine if this correlation was indeed given by a distinct differentiation state of the CD8low subpopulation, we next compared the expression of known CD8 differentiation markers in these cells with the CD8high subpopulation. Besides CD44 and CD62L, we evaluated the expression of other CD8 differentiation related molecules, the transcription factor EOMES[50], the IL-7 receptor CD127[51], and the terminal differentiation marker KLRG1[52] (Figures 1G and 1H). At the CD8high subpopulation, most of measured markers showed no significant differences to the exception of CD44, which was highly expressed in dKO CD8high cells. However, CD8low cells from the N2KO/N1Het and dKO mice were characterized by complete downregulation of CD62L and CD127, intermediate CD44 expression and reduction of EOMES when compared to CRE controls, although they showed no differences in KLRG1 levels. Consequently, we believe that cells in the CD8low population observed in the N2KO/N1Het and dKO mice were more differentiated than those seen in control mice. Phenotypically, marker expression of these cells defines them as short-lived and terminal effectors [53, 54]. Moreover, reduction in the expression of the coreceptor CD8 can be indicative of diverse situations, including transient and chronic responses to antigen, cytokine stimulation and STAT3 signaling dysregulation [55–58]. Hence, the combined deletion of PTTN1 and PTPN2 in CD8+ T cells, resulted in an increased in vivo differentiation into terminal effector cells with preferential accumulation in the spleen.

### PTPN1 deficiency potentiates pre-existent functional enhancement in PTPN2 KO CD8 T cells

To understand the functional capabilities of the CD8 T cells with combined deficiency of PTPN2 and PTPN1, we breed our colony of conditional mutant mice with OT-1 transgenic mice. OT-1 mice express a TCR with affinity for an epitope of chicken ovalbumin (OVA 257-64), recognized in the context of the MHC Class I haplotype H2-Kb expressed on C57BL/6 mice[59]. Naïve splenic CD8+ T cells from the different genotypes obtained and were then stimulated and expanded in vitro by receptor-crosslinking with antibodies for CD3 and CD28, and recombinant murine IL-2 to generate activated OT-1 specific CD8 T cells. We used naïve CD8 T cells to abrogate the potential biases originating from different frequencies of memory populations. Expanded cells conserved expression of the transgenic TCR α and β chains (Figure S2A). At day 4 post stimulation, expanded CD8 T cells were challenged with the OVA expressing thymoma cell line E.G7 in 5-hour in vitro cytotoxicity assays (Figures 2A, 2B, and S2B). In absence of a “third signal”, which is given by proinflammatory cytokines such as IL-12 and Type I interferons, naïve CD8 T cells will differentiate into activated cells with high memory forming potential and low effector capacities [60]. As expected, CRE controls followed such behavior, showing low cytotoxic potential (Figures 2A and 2B) and a favored differentiation towards Tcm cells (Figures 2C and 2D), a phenotype unaffected by the absence of PTPN1 expression (Figures 2A and 2B). Nonetheless, CD8 T cells with deleted expression of PTPN2 showed skewed differentiation to an effector phenotype characterized by high cytotoxic potential (Figures 2A, 2B, and S2B), and reduction of CD62L and CD44 expression (Figures 2C and 2D). The enhanced effector differentiation seen in the PTPN2 deficient cells is in accordance with previous reports [4, 5]. Despite that PTPN1 deficient CD8 T cells did not show any functional gain in cytotoxicity, when we analysed cells with hemi or complete deletion of PTPN1 on a PTPN2 deficient background, N2KO/N1Het and dKO genotypes respectively, the skewing towards an effector phenotype was even more potentiated (Figures 2A-D). Hence, absence of PTPN2 acted as a proinflammatory third signal that was further enhanced by the reduced activity of PTP1B, favoring differentiation towards an effector phenotype, while single deletion of PTPN1 was apparently uneventful.

**Figure 2.**
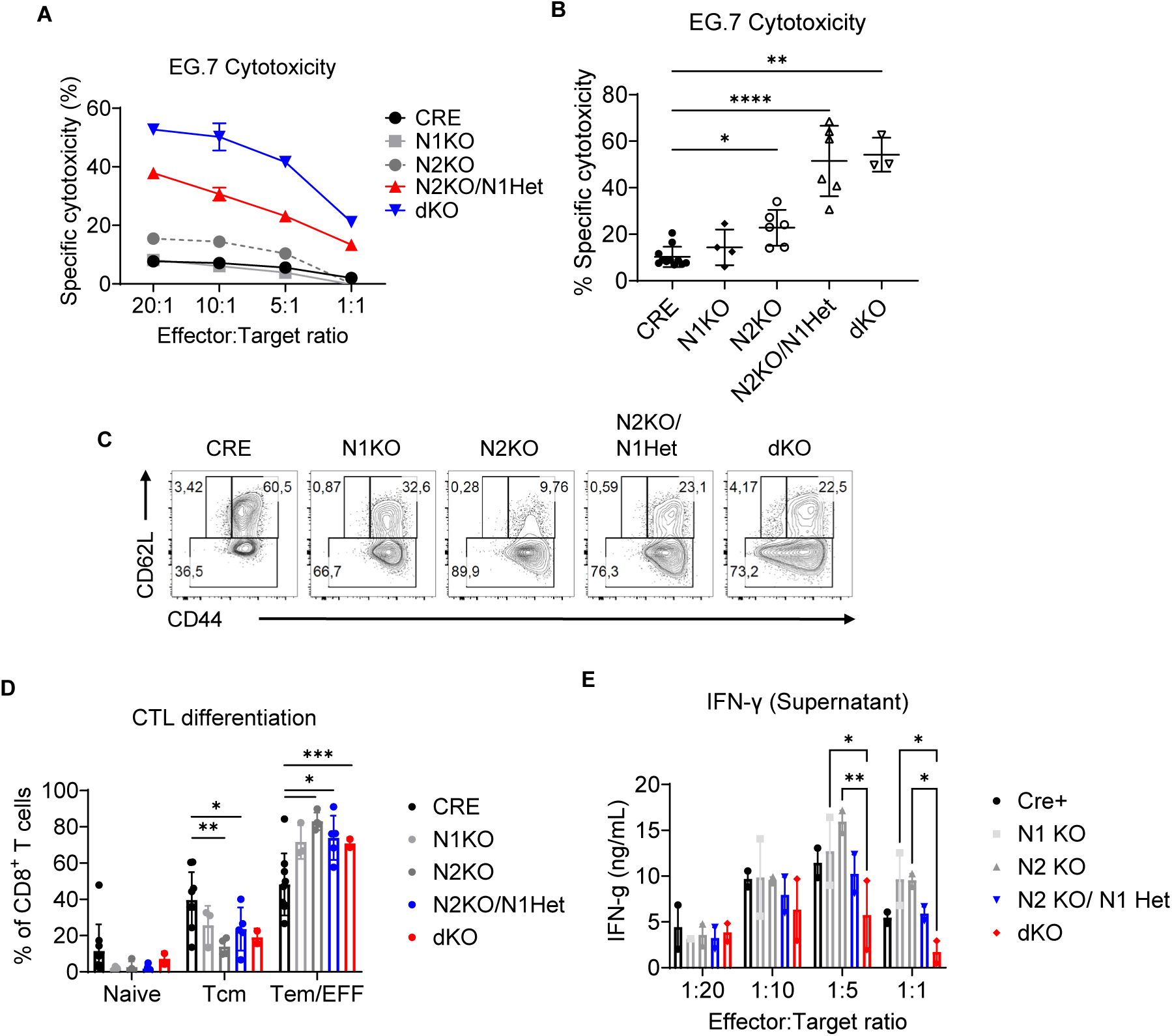
Combined deletion of PTPN2 and PTPN1 in CD8 T cells induce potent proinflammatory signals facilitating effector differentiation. Naïve CD8 T cells from the spleen of mice with different *Ptpn1* and/or *Ptpn2* deletions, expressing E8i-CRE and the OT-1 TCR, were activated/expanded with αCD3/αCD28 antibodies and IL-2. A) Cells at day 4 post stimulation were tested in vitro for specific cytotoxicity against the OVA-expressing thymoma E.G7. B) Statistical representation of several experiments. C) ELISA quantification of mouse IFN-γ in supernatants from (A). D) Flow cytometry plots of CD62L and CD44 expression from cells in (A) gated in singlet/ live events. E) Bar representation of the percentages of different populations from several experiments from (D). (All data representative of at least three different biological replicates; bars represent +/- S.E.M.; non-parametric Kruskal-Wallis test (B) or Mixed-effects analysis (C, E); P values *<0.05, **<0.005, ***<0.0005, ****<0.00005).

Secretion of proinflammatory cytokines, such as IFN-γ, is also a characteristic of effector CD8 T cells. To address the ability of mutant cells to produce IFN-γ, we performed ELISA in the supernatants collected from the cell cytotoxicity in vitro assays. We detected an increase in the secretion of IFN-γ in PTPN2 and PTPN1 single deficient cells at low effector-to-target ratios which was stronger in the PTPN2 KO cells (Figure 2E). However, no substantial changes were seen in the N2KO/N1Het and dKO T cells for most effector to target ratios, with the 1:5 and 1:1 ratios even showing lower IFN-γ production in dKOs (Figure 2E). Although secretion of IFN-γ and cytotoxic activity are both characteristic of effector CD8T cells, our results suggest that these functions are independently regulated. Moreover, while the shift towards an effector phenotype, defined by the loss of CD62L and CD44 expression, could explain the increased cytotoxic activity of N2KO, N2KO/N1Het and dKO CD8 T cells, we did not find a direct correlation between these two readouts (Figure S2C). Taking all together, the combined reduction of PTPN2 and PTPN1 activity resulted in the promotion of an effector phenotype and functionality, most likely as result of boosting diverse proinflammatory signals, and not solely IFN-γ.

### Signaling through the JAK/STAT pathway is largely dysregulated in PTPN1 and PTPN2 deficient cells

PTPN1 and PTPN2 are well known regulators of JAK/STAT signals. Proinflammatory cytokines “third signals” that induce effector differentiation, such as type I and 2 interferons, and IL-12, are transduced largely through this pathway. Hence, we sought to evaluate the c-terminal tyrosine phosphorylation of STAT proteins in these cells after proinflammatory cytokine stimulation. Type I interferons have a critical role in CD8 T cell acquisition of effector and memory functions [60, 61] and are known to signal through the phosphorylation of different STAT proteins in a context dependent manner [62]. To investigate how the combined absence of these phosphatases modifies STAT transduced type I interferon signals, we stimulated day 4 post activation CD8 T cells carrying different genotypes with IFN-β for 10 and 30 minutes. In our experiments, CRE control cells responded to recombinant murine IFN-β by increasing phosphorylation of these tyrosine residues in STATs 1, 2, 3, 4, and 5a (Figure 3A) revealing the complexity of type I interferon signals in CD8 cells. No increase in the phosphorylation of STAT6 was observed either in CRE control cells or in any of the genotypes analyzed (Figure 3B). The deficiency of either of these phosphatases resulted in the increased phosphorylation of STATs 1, 3, and 5, while phosphorylation levels of STAT4 seemed unaffected. A similar responsive pattern was observed in cells from N2KO/N1Het confirming that these cells are more sensitive to stimulation with type I interferons. Conversely, dKO cells were highly hyporesponsive with decreased phosphorylation of STATS 1, 3, and 5. Although the ratio of phosphorylated STAT4 to total protein was conserved, there was consistent decrease in the total abundance of STAT4 (Figure 3B).

**Figure 3.**
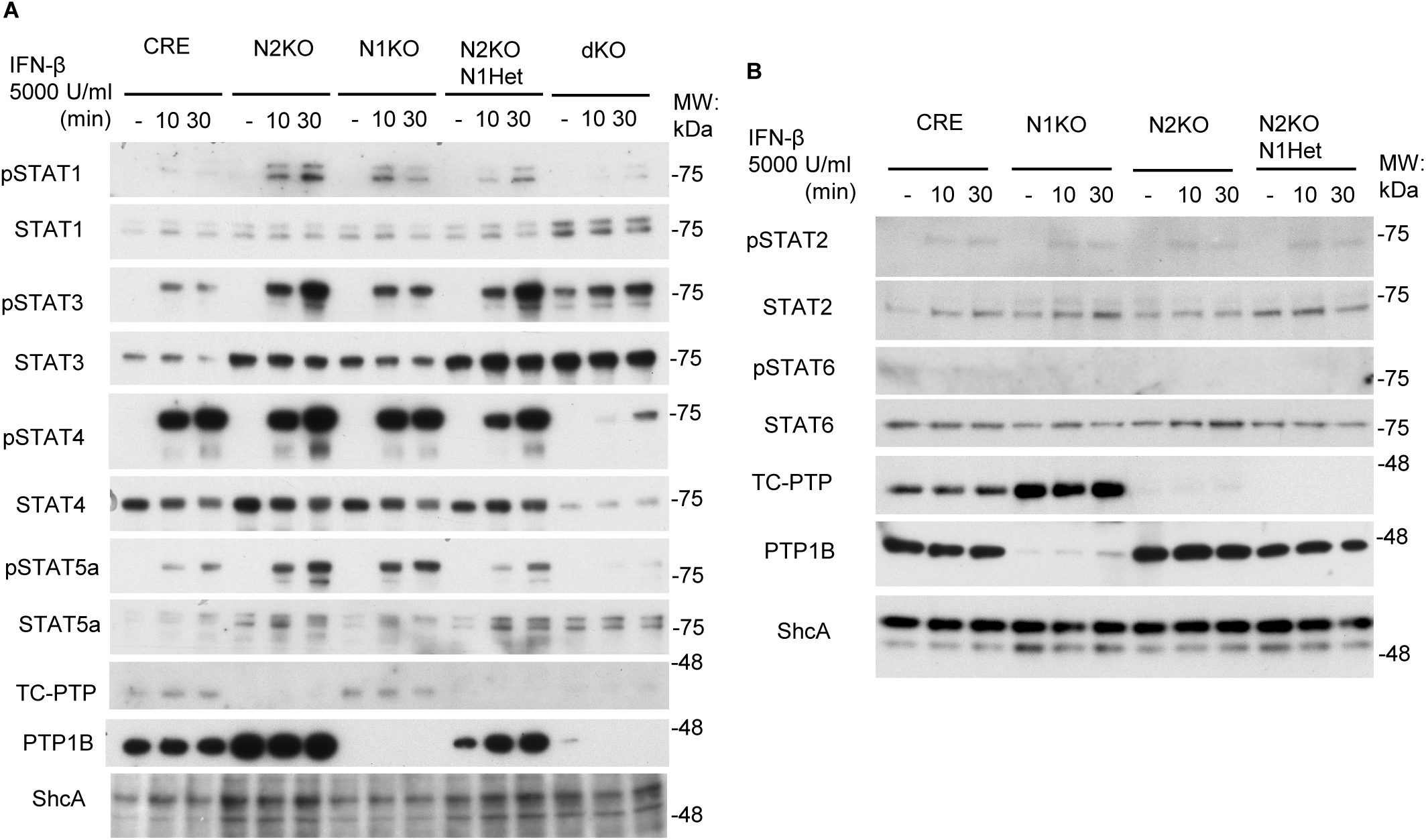
Reduction of PTPN2/PTPN1 activity induces global changes of STAT expression and phosphorylation in CD8 T cells. A) and B) Immunoblotting of protein lysates from day 4 post activation/expansion CD8 T cells, stimulated with 5000 Units of recombinant murine interferon β (rmIFN-β) for the indicated times. Time 0 controls correspond to cells without the addition of the cytokine. (Representative of at least 3 independent experiments).

CD8 T cells can also be responsive to INF-γ, which enhances their motility and cytotoxic capacity[63], however, CD8 T cells expanded in vitro do not respond to IFN-γ [64]. We reasoned then that the increase in the effector differentiation and cytolytic activity could arise from an autocrine loop involving IFN-γ in cells deficient of PTPN1 and/or PTPN2 by gaining sensitivity to the cytokine. Nevertheless, deficiency of PTPN1 and/or PTPN2 did not result in increased tyrosine phosphorylation of STAT1 or STAT3 upon IFN-γ stimulation in vitro (Figure S3).

Regardless of cytokine stimulation, resting cells revealed important differences in the regulation of the expression and phosphorylation of STATs in the different genotypes analysed. The largest discrepancy was observed in dKO cells which showed increased STAT 1 and 3 expression, in opposition to the observed STAT4 downregulation (Figure 3A and 3B). Moreover, we noticed a large increase in base-level phosphorylation of STAT3 in these cells (Figure 3A, 3B and S3), which was also observed at lower extent in N2KO and N2KO/N1Het cells (Figure 3A, 3B and S3).

As expected, reduced expression of PTPN1 and PTPN2 phosphatases resulted in the dysregulation of STAT tyrosine phosphorylation at both, baseline and after stimulation with IFN-β. Most importantly, basal STAT3 phosphorylation correlated with the acquisition of the effector phenotype observed previously.

### L598, a dual PTPN1/PTPN2 inhibitor, recapitulates genetic deficiency

The potentiation of effector functions observed on CD8 T cells after combined inhibition of PTPN1 and PTPN2 can reverse inhibitory cues in immunosuppressive microenvironments, such as tumors [7, 8]. To gain mechanistical understanding, we treated in vitro CD8 T cells with L598, a small, competitive, orthosteric and soluble compound that can inhibit PTPN2 and PTPN1 with equivalent IC50 [26, 65, 66]. To evaluate its ability to promote an effector phenotype as seen with other dual inhibitors, we activated splenic naïve CD8 T cells from OT-1mice following our in vitro protocol in presence of various concentrations (2, 5, and 10 µM) of L598 that were maintained during the differentiation/expansion process. We tested the treated CD8 T cells in cytotoxic assays against E.G7 targets (Figures 4A and 4B). We found that treatment with the compound significantly increased the cytotoxic activity of treated CD8 T cells against targets at all concentrations reaching its highest at 5µM. Nevertheless, we did not observe significant differences between 5µM and 10µM, uncovering a plateau concentration for the effects of the inhibitor. The enhancement of cytotoxicity seen in L598 treated cells was independent of the method used to produce the ex vivo CTL, as we were able to detect a similar increase in the cytolytic activity when these cells were generated by stimulation with the cognate peptide and APCs (Figure S4A). As cytokine secretion was also affected by the reduced activity of PTPN2 and PTPN1, we quantified IFN-γ secretion in the supernatant of these experiments by ELISA (Figure 4C). We observed a secretion pattern like the one seen on the genetically deficient cells, IFN-γ secretion was higher in the lower effector-to-target ratios, suggesting that low levels of stimulation favor proinflammatory cytokine secretion like what we had observed in cells with genetic deletion of these phosphatases.

**Figure 4.**
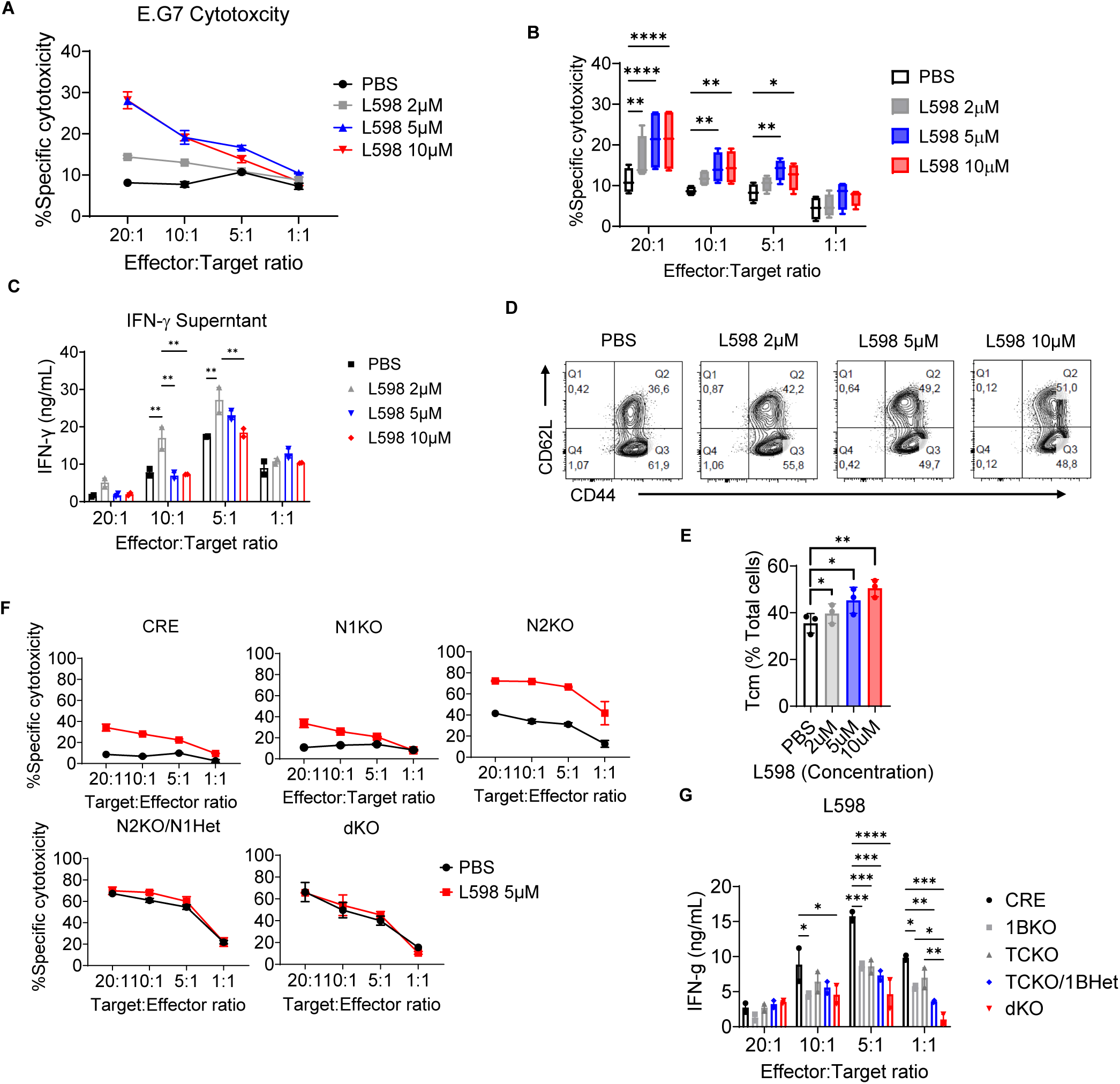
Pharmacological inhibition of PTPN2 and PTPN1 results in enhanced cytotoxicity and effector differentiation. CD8 T cells from OT-1 mice were activated/expanded in the presence of different concentrations of PTPN2/PTPN1 inhibitor L598 for four days. A) Specific cytotoxicity against OVA expressing thymoma E.G7 cells. B) Statistical representation of several experiments. (n=4) C) ELISA quantification of IFN-γ in supernatants from (A). D) and E) CD8 T cell differentiation determined by flow cytometric analysis of CD62L and CD44 expression in cells from (A). D) Flow cytometry plots of cells gated in single/live events. E) Tcm population percentages from different animals. (n=3) F) Specific cytotoxicity against E.G7 cells of CD8 T cells from animals with different PTPN2 and PTPN1 genotypes, additionally treated with L598 during activation/expansion. G) ELISA quantification of IFN-γ in supernatants from (F) (Bars represent +/- S.E.M.; One-way (E) or Two-way (B, C, G) ANOVA; P values *<0.05, **<0.005, ***<0.0005, ****<0.00005).

Next, we evaluated by flow cytometric analysis how the differentiation state of CD8 T cells was altered by the treatment with L598 (Figures 4D and E). Despite the functional enhancement observed with 5 and 10 µM, the expression of CD62L and CD44 drifted towards a modest but significant gradual increase of the Tcm population (CD62L high, CD44 high events; Q2) and correlated with the inhibitor concentration used (Figure 4D and E). Hence, treatment with L598 either potentiate the function of effector differentiated cells or confers effector characteristics to cells phenotypically characterized as Tcm. Moreover, when we activated CD8 T cells with the cognate peptide, differentiation instead drifted towards an increase in the proportion of effector/Effector memory cells when cells were in the presence of L598 (Figure S4B). In accordance, their cytotoxicity capacity was increased at least twice that observed in untreated cells (Figure S4A).

To dissect the role that pharmacological inhibition of each phosphatase plays on the onset of an effector phenotype, we treated CD8 T cells from our colony of conditional PTPN1 and/or PTPN2 mutants with 5µM L598 during activation and expansion and we tested these cells for specific cytotoxicity against E.G7 cells and IFN-γ secretion (Figure 4F). As seen in WT OT-1 CTLs, CRE controls increased their capacity to lyse target cells by about a fold. In the case of PTPN1 KO cells, cytotoxicity followed a pattern without significant differences to CRE controls either when treated or not with the inhibitor, suggesting a primary requirement of PTPN2 inhibition for the enhanced cytolytic activity observed in CD8 cells. However, after treatment with the inhibitor, PTPN2 KO cells still increased their cytolytic activity twice more than when compared to non-treated PTPN2 KO cells resembling the levels observed N2KO/N1Het and dKO cells. Hence, inhibition of PTPN1 with 5µM L598 had a comparable effect to the observations in the PTPN1 hemizygous cells. Moreover, the inhibitor did not influence either N2KO/N1Het or dKO CTLs cytotoxic capacities suggesting that, in both genotypes, cells are brought to their maximum effector capabilities (Figure 4F). In genetic deficient cells, IFN-γ secretion was even more reduced as result of the treatment with L598 when compared to L598 treated CRE controls (Figure 4G), supporting the existence of independent regulatory mechanisms for cytotoxicity and the secretion of IFN-γ.

### L598 also enhances human T cell functions

We tested, as well, the capacity of L598 to ameliorate the functionality of human T cells. To do so, we enriched T cells from healthy donors’ PBMCs and stimulated them in vitro with anti-CD3 and anti-CD28 for 7 days in the presence of increasing amounts of L598. On day 7 cells were harvested and analyzed by surface and intracellular flow cytometry for the pro-inflammatory cytokines IFN-γ and TNF-α, as for the differentiation markers CD45RO and CD62L defining effector and memory populations. In vitro treatment of human T cells with L598 led to an increase in the production and accumulation of both IFN-γ and TNF-α in both CD4 and CD8 T cells (Figures 5A-D). In all cases, the increase reached its maximum with 2µM treatment and was maintained at a similar extent when cells were treated with increasing concentrations. The increase in cytokine production was around 50 to 70% for IFN-γ (Figure 5B) and 30 to 40% for TNF-α (Figure 5C), in both CD4 and CD8 T cells. Besides increased proinflammatory cytokine production, we also observed that treatment with L598 skewed T cell differentiation towards a central memory phenotype (CD45RO high, CD62L high) in both CD4 and CD8 T cells (Figure 5D). This increase was more pronounced in CD8 T cells and seemed to correlate to the concentration of L598 used. We did not find any preference for improving CD4 or CD8 T cell proliferation as both cell types were similarly represented at the different concentrations used of L598 (Figure S5).

**Figure 5.**
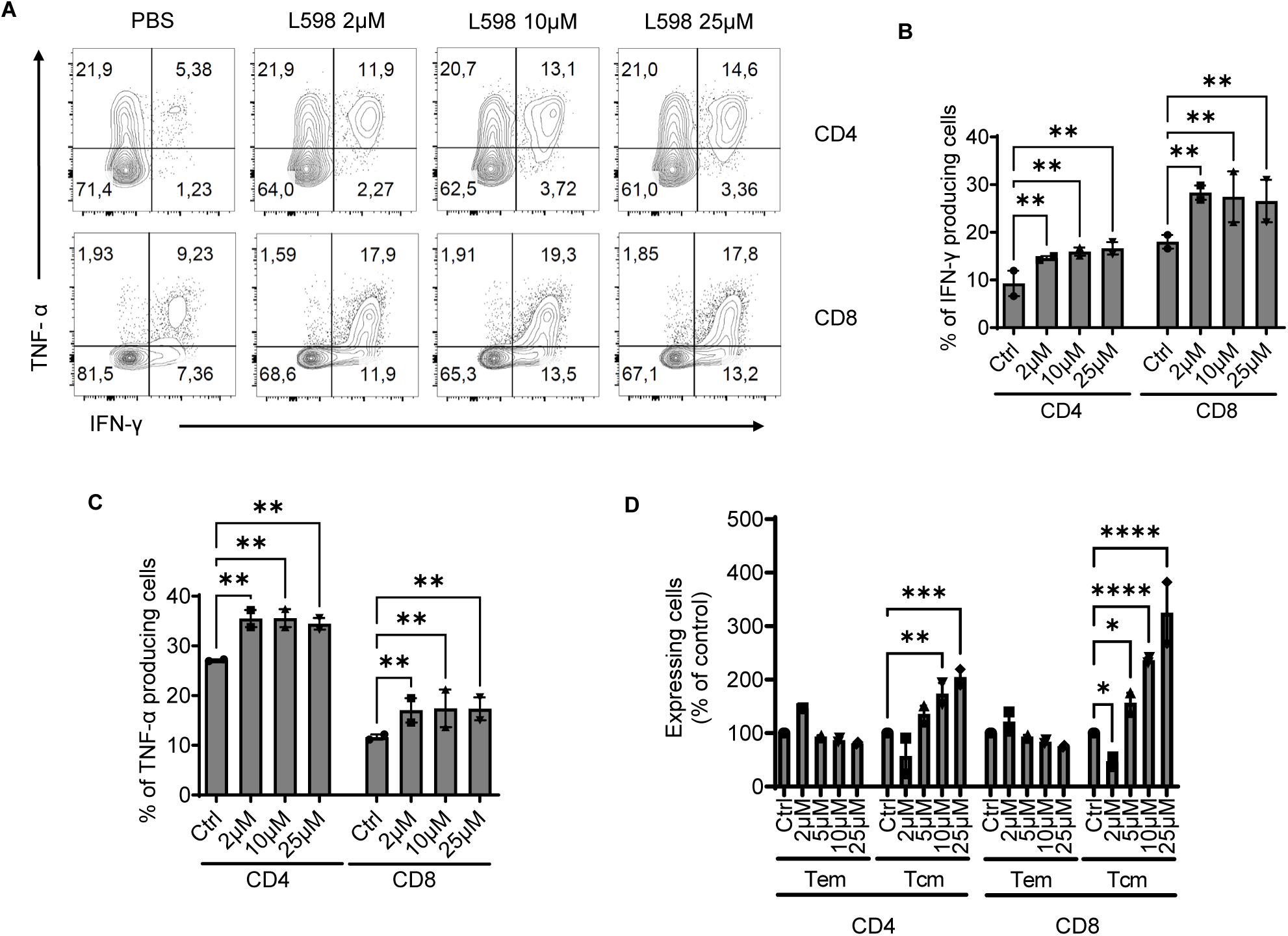
L598 treatment enhances effector differentiation in human T cells. A) Purified T cells from human donors were activated by αCD3/αCD28 and analyzed on day 7 for IFN-γ and TNF-α by flow cytometric analysis. Plots of cells gated in CD4 and CD8 populations distributed by IFN-γ and TNF-α expression. B) and C) Bar graphs showing the statistical representation of 2 biological replicates. D) Bar graphs of cell populations sorted by expression of CD45RO and CD62L defining Tem (CD45RO^high^; CD62L^low^) and Tcm (CD45RO^high^; CD62L^high^) phenotypes. I) Percentual contribution of CD4 and CD8 T cells of in vitro human T cell cultures at day 7. (Bars represent +/- S.E.M.; Two-way ANOVA; P values *<0.05, **<0.005, ***<0.0005, ****<0.00005).

### Transcriptomic analyses suggest functional dominance of PTPN2 over PTPN1 in CD 8 T cells

Combined deficiency of PTPN1 and PTPN2 resulted in synergistic signals driving the differentiation CD8 T cells into effectors as result, in part, of changes in the expression and activation of STAT transcription factors, either at baseline or after cytokine stimulation. However, both phosphatases are known regulators of other pathways relevant in T cells [28]. To understand the extent of the transcriptional changes promoted by the deficiency of PTPN1 and PTPN2, we performed bulk mRNA sequencing on day 4 stimulated naïve CD8 T cells derived from the different genotypes. We aimed to identify the differentially regulated transcripts by comparative RNAseq (Figures 6 and S6). Unsupervised principal component analysis (PCA) revealed that the PTPN1KO and PTPN2 KO samples drifted from CRE controls following a divergent pattern (Figure 6A), strongly suggesting that both phosphatases target, at least partially, non-redundant pathways. The drift observed in the PCA was also reflected in the number of significant differentially expressed (DE) (Figure S6A). Interestingly, while PTPN2 KO cells showed a stronger effector phenotype than PTPN1 KO cells, the amount of DE genes was relatively smaller. We additionally analyzed the transcriptional changes induced by the dual inhibitor L598 in day 4 differentiated/expanded CD8 T cells from WT C57BL/6 mice treated or not with 5µM L598. Treatment of CD8 T cells with L598 resulted in a large number of DE genes, similar to that observed in the N1KO and dKO genotypes (Figure S6B). Organizing the data in Venn diagrams (Figure 6B) revealed that 66% of the upregulated and 64% of the downregulated genes after L598 treatment corresponded to genes observed in the genetically deficient cells analysed, reassuring that transcriptional modifications are the result of PTPN1 and PTPN2 inhibition. The Venn diagrams also revealed that all but one of the DE genes identified in the PTPN2 KO cells were also significant in the N2KO/N1Het background, and still a large proportion of those, nearly 90% of downregulated and 65% of upregulated, were identified in dKO cells. On the contrary, PTPN1 KO cells despite having the strongest transcriptional drift from CRE controls (Figure 6A), did not represented genes observed in other genotypes, which was also evident when plotting the data as a heatmap (Figure 6C). Roughly 83% of downregulated and 82% of upregulated genes were uniquely found in the PTPN1 KO genotype. While comparing both single KOs, just a very small proportion of genes found in common between the PTPN2 KO and PTPN1 KO cells, 17 of the downregulated and 22 of the upregulated genes, corresponding to 11.6% and 13.3% of total PTPN2 KO DE genes, highlighting functional divergence between both phosphatases.

**Figure 6.**
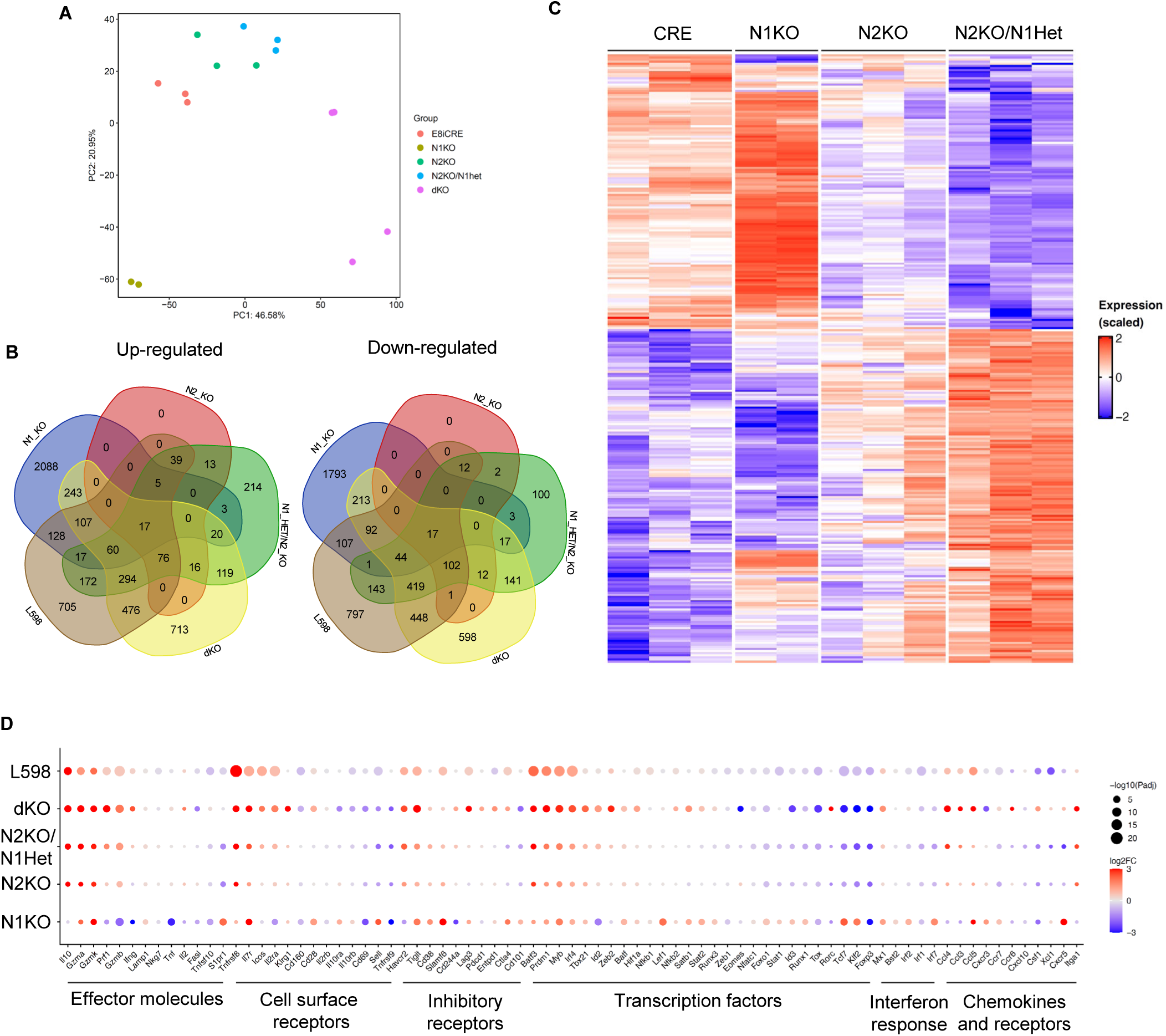
Bulk mRNA analysis of CD8 T cells deficient in PTPN2 and/or PTPN1 or treated with L598. A) Principal component analysis (PCA) of bulk mRNA sequencing samples obtained from day post stimulation/expansion from different genotypes analysed. B) Venn diagrams illustrating significant differentially expressed genes in samples from (A) and those form L598 treated vs untreated WT C57BL/6 mice (FDR 5%). C) Heatmap of genes differentially expressed in the genotypes analyzed and organized as opposing expression between N2KO/N1Het CD8 T cells and CRE controls. Overexpressed genes showed a gradual red shift, and under-expressed genes a gradual blue shift. D) Dot plot corresponding to a selected list of genes relevant to CD8 T cells activation/differentiation. Overexpressed genes showed a gradual red shift, and under-expressed genes a gradual blue shift.

To better analyse the large amount of data generated, we used a biased approach to emphasize a transcriptomic signature relevant to CD8 T cell biology (Figure 6D). As our previous data suggested, PTPN2 deficiency corresponded to the skew towards an effector phenotype characterized by an increase in the expression of messengers for granzymes and perforin, the reduction of those related to secondary immune organs migration, such as CD62L and CCR7, mild expression of inhibitory receptors such as PD1, Tigit and Tim-3, and increase of CCL3, 4 and 5 chemokines. Upon concomitant heterozygous deletion of PTPN1 (PTPN2 KO/N1Het cells), this effector signature observed under PTPN2 deficiency was not only maintained but even reinforced, a finding that was further extended in dKO cells. In opposition, we observed the same tendency seen with the Venn diagrams, where PTPN1 deficient cells showed a very contrasting transcriptomic signature when compared to all other genotypes analyzed. In PTPN1 KO cells we observed reduction of the transcripts for effector molecules such as Granzyme B, Perforin and TNF, upregulation of the those for the CD127, CD62L, CXCR5, and CD28 surface receptors, as of the transcription factors Tcf1 and Id3, corresponding to a T central memory signature [67]. Accordingly with the observed effector phenotype, the transcriptome of CD8 T cells treated with L598 highly resembled that of the N2KO and N2KO/N1Het yet confirming that the mechanisms improving the effector qualities of cells treated with L598 are given by similar changes observed in the deficient genetic cells. Although the transcriptome of dKO cells correlated to that of single PTPN2 KO cells, these cells also expressed higher levels of receptors associated with terminal differentiation, such as Tim-3, Tigit, Lag3, Entpd1, and PD-1. Also, we observed in dKO cells the ectopic expression of the Th17-related transcription factor Rorc suggesting that transcriptional dysregulation in these cells could result in aberrant differentiation transition to Th17-like phenotype, a phenomenon observed in CD4 Treg cells with PTPN2 loss of function mutations [68].

### CD8 T cells with combined deficiency of PTPN1 and PTPN2 represent an alternative effector differentiation route

The resulting transcriptomes are a consequence of dysregulation of numerous pathways which components are substrates of PTPN1 and PTPN2. Gene Set Enrichment Analysis (GSEA) allows to compare to existing transcriptomic signature to curated signatures previously obtained. The Hallmark gene sets collection provide the recollection of redundant genes well-stablished in particular biological states [69]. We found that transcriptomic dysregulation caused by the absence of PTPN1 scored in a larger number of Gene Sets than other genotypes (Figures 7A and S7A). Of note, pathways well known for transducing through the JAK/STAT pathway such as IL-2, type 1 and type 2 interferons were significant in PTPN1 KO cells while not affected in other genotypes (Figure 6A). Contrary to expectations, PTPN1 KO cells showed a positive correlation with type 1 and type 2 interferon signaling, while other genotypes were not significant or even showed a small negative correlation (N2KO/N1Het). These results suggest that type 1 INF signals are not responsible for the skewing towards effector differentiation seen in genotypes where PTPN2 signals are dominant (N2KO, N2KO/N1Het and dKO). Only other STAT related pathway scored significantly, IL-2/STAT5, however it correlated negatively in PTPN1 KO cells (Figure 7A). This same genotype showed downregulation of the TGF-β and TNF-α pathways (Figure S7A). Besides cytokine pathways, other two pathways of the Hallmark gene sets scored significantly, Myc and P53 (Figure 7A). Myc regulation is known to favor protein synthesis, mitochondrial biogenesis, and activation of glycolytic pathways in CD8 T cells [70, 71]. In these cells, P53 collaborate in the maintenance of this phenotype through a feedback mechanism involving Myc [72]. While these data fits with the differentiation towards an effector phenotype, is worth to point out that prolonged activation of this axis is associated with exhaustion onset [73]. Nevertheless, although N1KO cells showed transcriptional changes indicating reduced glycolysis, no significant changes related to metabolic regulation of oxidative phosphorylation or glycolysis were observed (Figure S7A).

**Figure 7.**
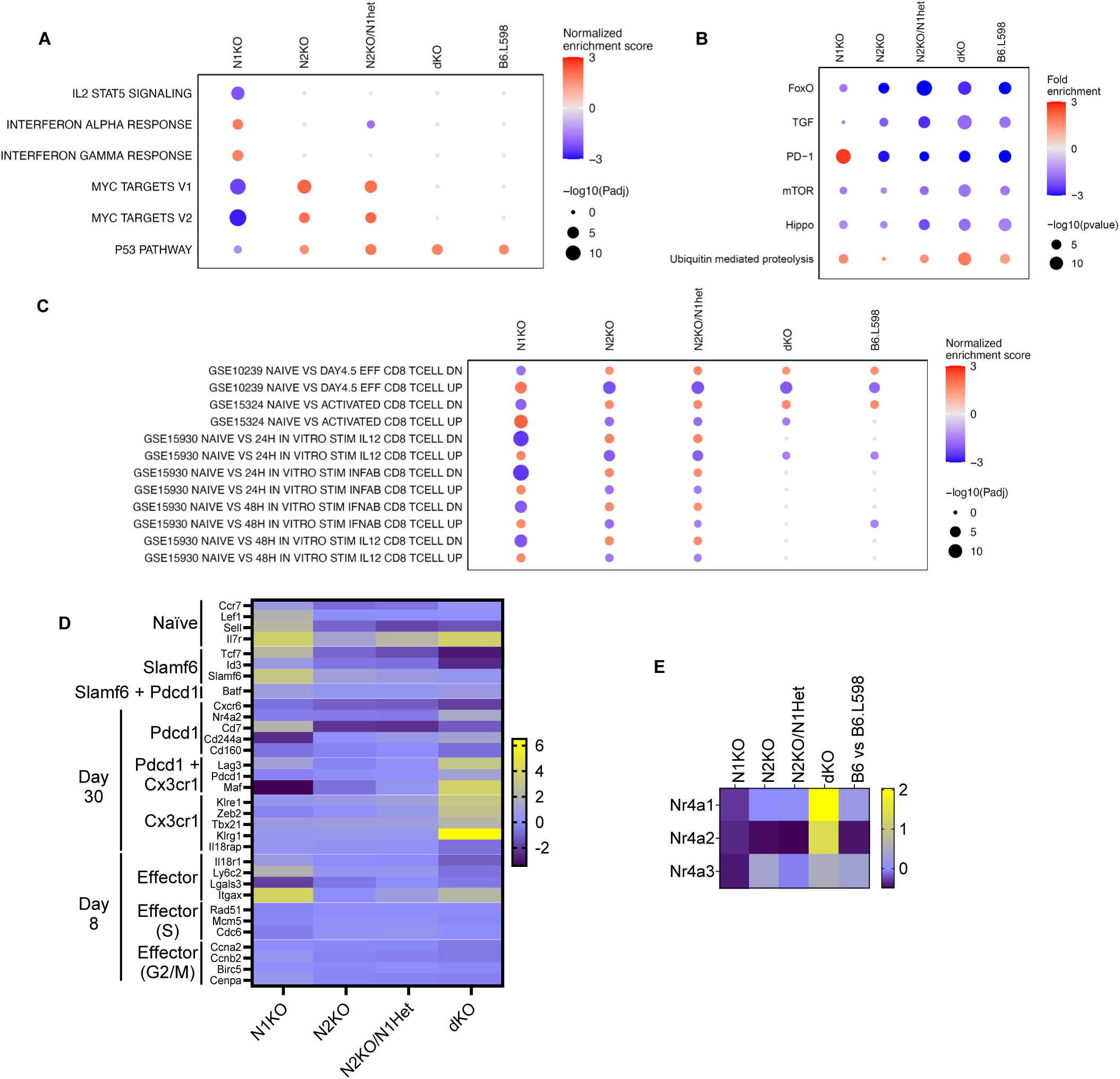
Analytic transcriptomics suggests that PTPN1 and PTPN2 inhibition promotes high functioning CD8 T cells terminal effector differentiation. A) Selected significantly scoring Hallmark collection gene sets from the transcriptomes of different PTPN1 and/or PTPN2 deficient CD8 T cells and L598 treated cells (additional significantly scoring gene sets can be found in the supplementary material Figure S6). B) Significantly scoring pathways after pathfindR analysis of same samples as in (A). C) GSEA analysis of samples in (A) transcriptomes obtained in similar analysis from (Sarkar et al. [51]: GSE10239, Yamada et al. [74]: GSE15324 and Agarwal et al.[59]: GSE15930.). D) Heat map of transcript expression in samples from (A) corresponding to those genes listed in Zander et al.[41] defining specific CD8 clusters. E) Heat map of the transcript expression for members of the Nr4a transcription factor family in samples from (A).high-functioning

Despite divergent transcriptomic signatures, PTPN1 and PTPN2 single deficiency can both confer effector advantages in vivo [2, 4–6]. To gain insight on the nature of the signals favored by the absence of either or both phosphatases, we performed pathway mapping analysis on the transcriptomes of these cells with pathfindR, a tool that takes into consideration subnetworks of protein-protein interaction [74], hence allowing us to predict the responsiveness to different stimulus (Figures 7B and S7B). We found a significant downregulation of components of the FoxO, mTOR and Hippo pathways, most likely as a feedback mechanism in response to the metabolic switch signals promoting glycolysis. Again, as a representation of the dichotomic nature of N1KO vs N2KO signals, N1KO cells scored positively to PD-1 pathway while the rest scored negatively, suggesting those genotypes dominated by PTPN2KO signals could show resistance to PDL 1 and 2 inhibition.

GSEA allowed us also to compare our data with previously generated CD8 transcriptomic data [52, 60, 75]. Contrary to expectations, we found that absence of PTPN1 reinforced the signature of CD8 cells activated in vivo after 4 days, while this signature was negatively correlated by those genotypes dominated by the PTPN2 deficiency and that of L598 treated cells (Figure 7C). Moreover, the same correlation was observed in cell stimulated in vitro in addition to the proinflammatory cytokines IL-12 or type 1 interferons (Figure 7C). Although it might seem counterintuitive taking into account our phenotypic and functional data, it is well understood now that CD8 differentiation does not follow a linear model, instead it is highly diverse and responds according to the type and length of the stimuli [76, 77]. Indeed, understanding of CD8 T cells differentiation has progress extensively in the later decade, accelerated by the acquisition of single cell transcriptomics. Zander et al. generated, by unsupervised clustering, a selected a set of representative genes that allows the determination of differentiation path of CD8 T cells after immunogenic stimulation in the LCMV model [42]. We cross-referenced their published gene list with the transcriptomic data obtained from the genetic deficient cell, allowing us to get an estimate of their lineage (Figure 7D). Deficiency of PTPN1 resulted in cells upregulating transcripts associated with naïve (*Lef1*, *Sell*, *IL7r*), effectors (*Ly6c2*, *Itgax*) and *SLAMf6* expressing self renewing progenitors (*SLAM6*, *Tcf7*), while PTPN2KO and PTPN2KO/1BHet cells did not showed a particularly reinforced transcriptomic signature. However, dKO cells did increased the expression of transcripts related to exhausted cells (*Lag3*, *Maf*, *Klre1*, *Zeb2* and *Klrg1*), more specifically related to the terminally differentiated CX3CR1 highly functional phenotype. Despite this result, it is important to remark that naïve dKO cell reach this transcriptomic signature only after 4 to 5 days post activation. Hence, total deletion of the activities of PTPN1 and PTPN2 in CD8 T cells could accelerate the onset of chronic or dysfunctional differentiation states. Indeed, dKO cells also showed increased upregulation of the three members of the transcription factor family NR4A (Figure 7E) in dKO cells over all the other genotypes analyzed, which is associated with the establishment of exhaustion in CD8 T cells [78]. However, no upregulation of the exhaustion-related transcription factor Tox was observed (Figure 6D). Taking our transcriptomic data together, we concluded that despite seen a dysregulation of STAT phosphorylation, the combined deficiency of PTPN1 and PTPN2 was not interpreted by CD8 T cells as an enhancement proinflammatory signals derived from interferons or IL-12 but correlated to the formation of highly functional terminal differentiated cells seen in chronic infectious diseases and cancer.

### Autocrine IL-10 is responsible for the gained effector differentiation

We found the IL-10 transcript to be one of the highest upregulated genes which also followed a gradient pattern that correlated to the gains in cytotoxicity observed in N2KO, N2KO/N1Het and dKO cells (Figures 6D and S8A). IL-10 is regarded as a regulatory cytokine, as systemically it induces tolerance [31, 79, 80] and supresses the activity of CD4 cells [30], however, it acts as a promoter of tumor immunity [33, 81] and enhances the activity of CD8 T cells[41]. Therefore, we decided to determine if the expression of the transcript was followed by production and secretion of the cytokine. To this end, we tested the supernatants from cytotoxic experiments for the presence of the IL-10 by ELISA (Figure 8A). Indeed, we found that N2KO cell secreted IL-10 at levels 3-4 times higher than control CRE cells, peaking at around 50 to 60 pg/ml. However, more strikingly, secretion of IL-10 by N2KO/N1Het and dKO cells were more than 20 to 60 times respectively to that seen in single N2KO cells. As a result, we concluded that PTPN2 deficiency promoted the differentiation to effector CD8 T cells which secrete IL-10, and concomitant deficiency, either partial or total, potentiated this effect. We also observed upregulation of the IL-10 transcript in CD8 cells after treatment with L598 (Figure S8B). To ascertain if the L598 inhibitor would also promote the secretion of IL-10 and if it is potentiated in a genetic deficient background, we tested the secretion of IL-10 in supernatants of those cytotoxic experiments (Figure 8B). We found that treatment with L598 indeed increases the secretion of IL-10 in CRE controls and potentiated the effect in cells with genetic deficiency of either or both phosphatases.

**Figure 8.**
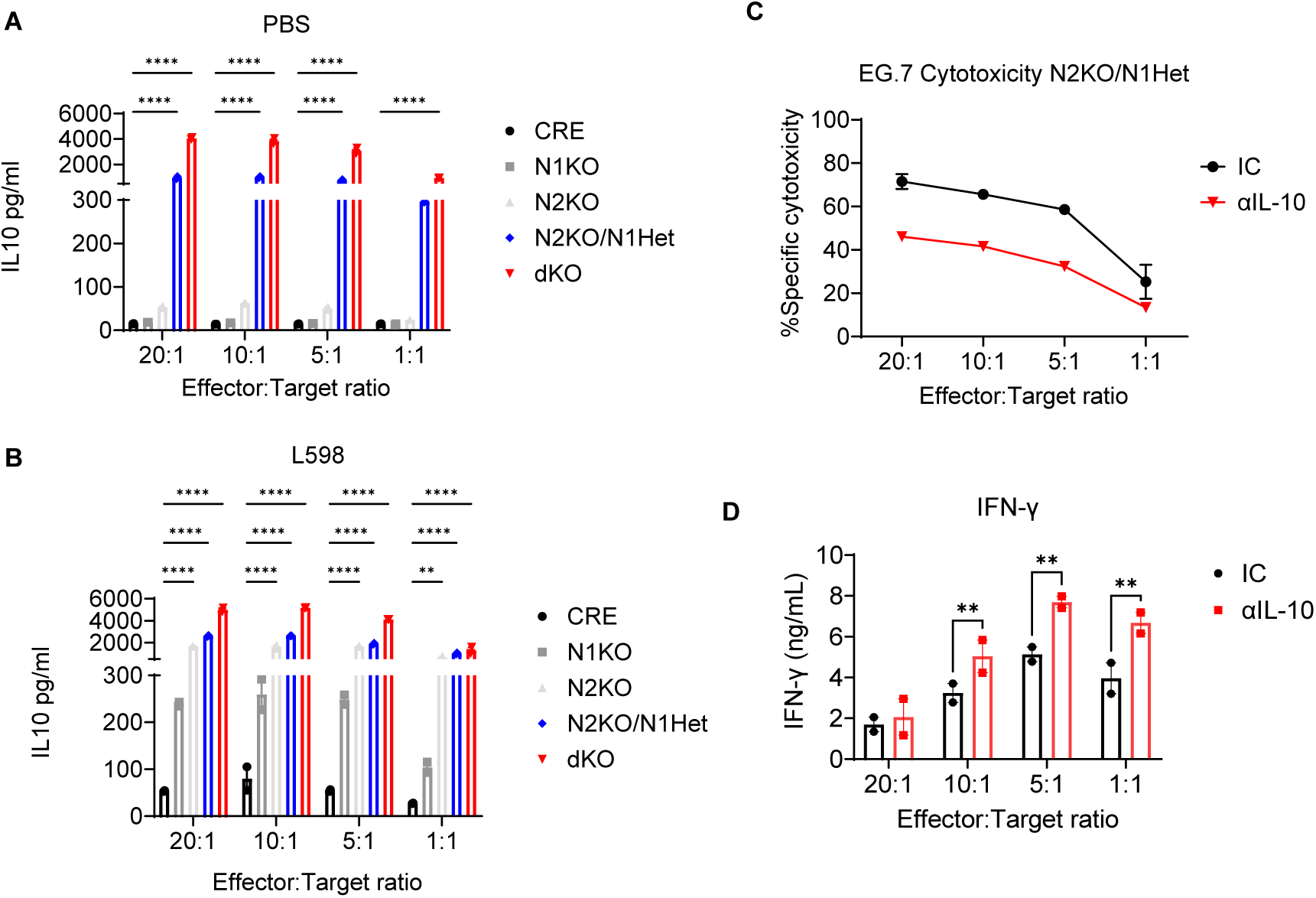
Autocrine stimulation by IL-10 is responsible for the functional gain in N2KO/N1Het CD8 T cells. Secretion of IL-10 from day 4 activate/expanded CD8 T cells carrying different TC-PTP and PTP1B mutations, A) without or B) after with 5µM L598. IL-10 concentrations were determined by ELISA from supernatants of experiments in Figure 4F. C) Specific cytotoxicity against OVA expressing E.G7 thymoma cells from day 4 N2KO/N1Het activated/expanded CD8 T cells incubated in the presence of an IL-10 neutralizing antibody or an isotype control (Representative of 3 experiments). D) ELISA quantification of mouse IFN-γ in supernatants from (C). (Bars represent +/- S.E.M.; Two-way ANOVA; P values *<0.05, **<0.005, ***<0.0005, ****<0.00005).

As IL-10 can potentiate the activity of CD8 T cells, we decided to evaluate the role of this cytokine in the effector differentiation of N2KO/N1Het naïve CD8 T cells. To this end we generated in vitro effector CD8 T cells from this genotype in the presence of a neutralizing IL-10 antibody or an isotype control. Upon 4 days of activation and expansion, we tested these cells for EG.7 cytotoxicity and secretion of INF-γ (Figure 8C and 8D). We observed a reduction of the cytotoxic capacity of these cells close to 50%, while significantly increased the secretion of IFN-γ opposing the phenotype given by the concomitant hemi-deletion of PTPN1 in a PTPN2 deficient background. Hence, by blocking the activity of IL-10 we reversed the phenotypes acquired by the combined deletion of PTPN2 and PTPN1.

### KQ791, a derivative of L598 synergizes with PD1 blockade to in a partially responding breast cancer model

PTPN1 and PTPN2 regulate pathways alternative to classic checkpoint targets as PD-1, CTLA-4, TIM-3 or LAG-3[28]. It stands to reason that combinational therapies targeting checkpoints and PTPN1/2 inhibition could result in enhanced tumor control. To test this hypothesis, we made use of a clinically tested (Trial IDs phase IA NCT02445911 and phase IB/II NCT02370043) derivative of L598 named KQ791. KQ791 exhibited higher potency than L598 in in vitro phosphatase assays displaying IC50 for both PTPN1 and PTPN2 3 times lower (Figure 9A). Similarly, KQ791 required a concentration 5 times lower during naïve CD8 T cell activation/expansion to achieve equivalent increase in biological activity, measured by specific cytotoxicity against E.G7 cells (Figure 9B). As preclinical and clinical data demonstrated high tolerability of KQ791 in animal models and humans, we decided to test its ability to inhibit tumor growth in vivo in the syngeneic breast cancer model EMT-6, known to respond partially to PD-1 blockade therapy (Figures 9 C-E). Despite observing a marginal and not significant reduction in the tumor growth of animals subjected to either treatment alone, KQ-791 or α-PD1, the combination of both inhibited the tumor growth (Figure 9C), significantly increased the survival (Figure 9D) and the number of mice reaching complete regression (CR), 7 in the double treatment group vs 2 in both single treatments and none in the control group (Figure 9E).

**Figure 9.**
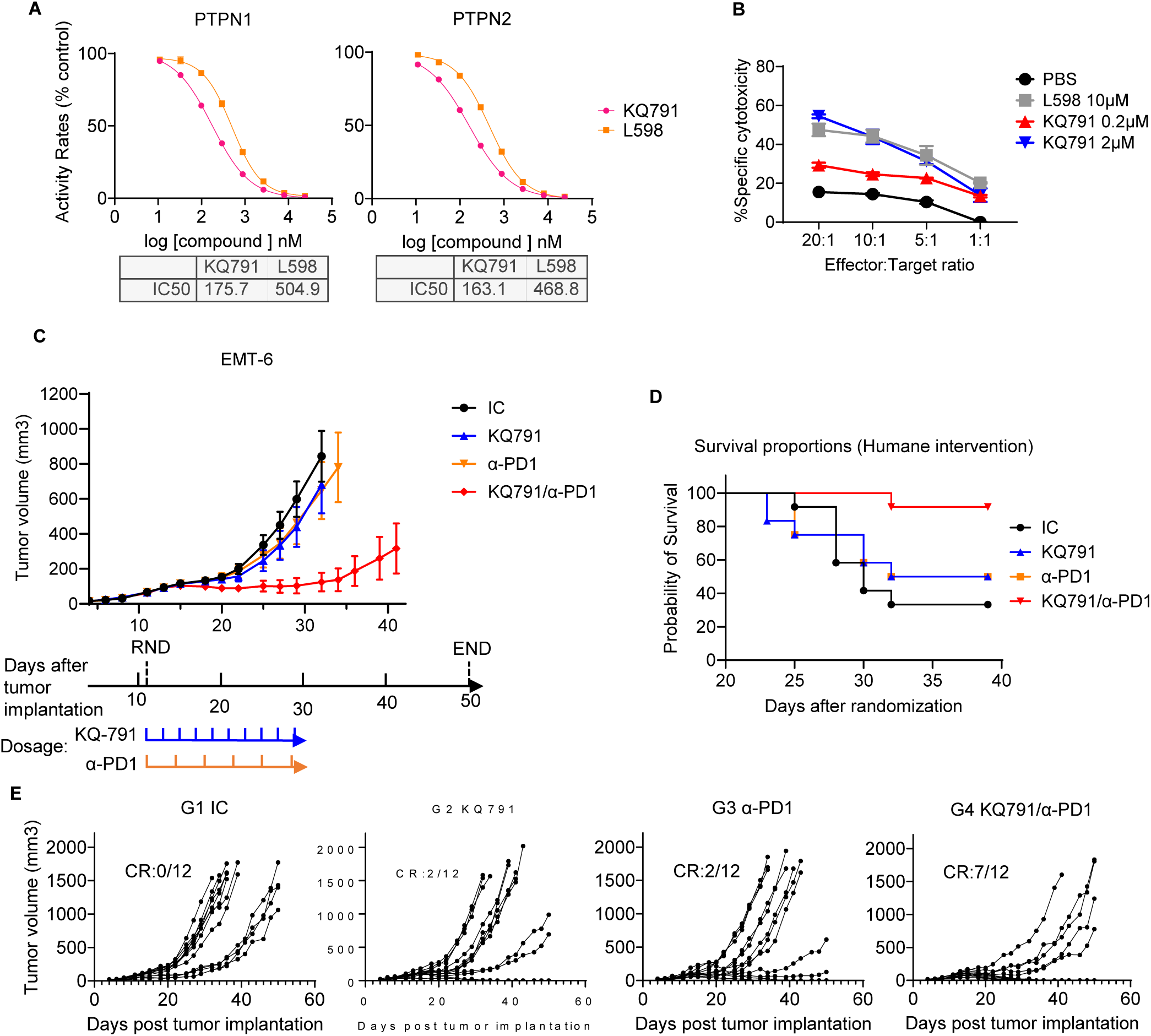
Pharmacological inhibition of PTPN1 and PTPN2 synergizes with anti-PD1 blockade therapy to restrict EMT-6 tumour growth in vivo. A) In vitro phosphatase activity of PTPN1 and PTPN2 catalytic domains in presence of increasing concentrations of L598 and KQ791. Values are the average IC50 from 3 separate experiments. B) Specific cytotoxicity against OVA expressing E.G7 thymoma cells from 4-day C57BL/6 CTLs treated with L598 or KQ791 at the concentrations specified. Representative of 3 independent experiments. C) Timeline of tumour volume after subcutaneous injection of 1.5 x 10^5^ EMT-6 breast cancer cells in BALB/c mice. Mice were subjected to treatment with an isotype control (IC), KQ791, anti-PD1 mAb (α-PD1) or both. (In all groups n=12) D) Kaplan-Meier survival analysis determined by humane end point intervention (See Materials and Methods). E) Timeline plots with the individual tumor growth curves, CR: number of mice reaching complete remission

## Discussion

Inhibition of PTPN1 and PTPN2 either individually or in combination has been proposed as a powerful immunotherapeutic target to enhance antitumoral immune responses [2, 3, 5, 6]. Moreover, reduction of PTPN2 activity in tumor cells facilitates interferon signals rendering them more susceptible to cytolysis of CD8 T cells[3], hence resulting in a win-win situation regarding cancer immunotherapy. The use of small molecule inhibitors targeting both enzymes had already demonstrated utility in mouse models [7] and is currently being explored in clinical trials by Calico and ABBIE (NCT04777994 and NCT04417465). Mechanistically consequences of the combined inhibition are not well known, however both phosphatases are highly homologous[1], and their known substrates belong to common families of phosphoproteins [14–16, 18–21, 23], prompting speculations about their specificity and redundancy. Similarly, development of orthosteric inhibitors that target specifically either of these phosphatases has proved difficult given the similarities between their catalytic domains [25–27], nevertheless, they demonstrate high specificity regarding other members of the PTP family. This dual specificity appears to confer them a particular advantage as immunotherapeutic agents, assuming non-redundant roles for PTPN1 and PTPN2. To better understand the mechanistic consequences of dual inhibition, here we explored the genetic deletion of PTPN1 and/or PTPN2 in mature naïve CD8 T cells, main cellular responders in tumor immunology, and compare them to the treatment with a dual pharmacological inhibitor.

We found that removal of PTPN2 was interpreted by CD8 T cells as a proinflammatory “third signal” favoring their differentiation into effector cells with increased cytotoxic and IFN-γ secretion capabilities. In contrast, deficiency of PTPN1, despite displaying a larger transcriptomic dysregulation, did not resulted in evident phenotypical or functional differences when compared to CRE controls. As result, our data agrees with the proposed non-redundant role of both phosphatases, where PTPN2 has evolved as a more immune related regulator while PTPN1 has a ubiquitous presence as a metabolic mediator. However, when PTPN1 was inhibited under a PTPN2 deficient background, either genetically or pharmacologically, it magnified the effector differentiation and the enhancement of cytotoxicity seen in the PTPN2 KO CD8 T cells. This reinforcement of the phenotypical changes given by the combined inhibition of PTPN2 and PTPN1 activity was also evident in the transcriptomic signature. Under a PTPN2 deficient background, upregulated and downregulated genes followed a gradual increase that correlated with the expression of PTPN1, WT>HET>KO, but drifted away from the PTPN1 KO signature. Interestingly, we found the transcriptomic signature induced by the dual PTPN1/PTPN2 inhibitor L598 followed that of the dKO, suggesting, as expected, it is a consequence of the specific inhibition of those phosphatases.

Our data supports the existence of an important house keeping role for these phosphatases, which appears to be redundant. When analysing either of the single deficient PTPN1 or PTPN2 CD8 T cells, they lacked a strong post thymic phenotype evidencing similar peripheral distribution and differentiated populations to those seen in CRE controls. Nonetheless, as consequence of double deficiency change were notorious, as naïve cells were found in scarce numbers with most peripheral CD8 cells identified as either as effector or memory cells. Hence the activity of either of them is required to maintain quiescence. Activated double deficient cells CD8 T cells then induced involution of the thymus around 10 weeks of age. This phenotype is observed in infections inducing cytokine storms, particularly those involving IFN-γ [82], encouraging us to speculate a similar mechanism in dKO mice. Interestingly, our previous observations in germline double KOs indicated that this condition was embryonically lethal [22], whereas presence of a copy of either PTPN1 or PTPN2 was sufficient to allow full-term gestation. Study of the embryos revealed that dead occurred around E9.5 and was characterized by hyperphosphorylation of STAT1, which was absent in any other genotype. Similarly, in the case of CD8 conditional mutants, presence of one copy of either PTPN1 or PTPN2 was sufficient to prevent the onset of thymic involution. Even though in the case of mice with PTPN2KO/PTPN1Het CD8 T cells where we found a similar accumulation of CD8low effectors in the periphery. Double deficient cells also displayed a transcriptome more extensively dysregulated and compatible in several with that of chronically stimulated cells, given by the higher expression of several inhibitory receptors (Tim3, Tigit and Lag3) and the exhaustion associated transcription factors of the Nr4a family, despite been stimulated only for 4 days. Still, these cells did not express larger quantities of the transcription factor Tox and were more compatible in their signature to terminal CX3CR1 effectors than PD-1 expressing exhausted cells and were highly functional. Similar observations have been done previously in vivo with single PTPN2 deficient cells, which reversed the onset of exhaustion [4].

Mechanistically, our results indicate that the combined inhibition of PTPN1 and PTPN2 resulted in the increment of baseline STAT3 c-terminal tyrosine phosphorylation which later we correlated to the secretion of IL-10. Moreover, previously we had found that PTPN1 deficiency in macrophages induced enhanced expression of IL-10 dependent regulatory genes such as IL-4Rα through STAT3, but concomitantly promoted aberrant STAT1 proinflammatory signals [39] suggesting a similar effect in CD8 T cells which leaned more clearly towards a proinflammatory phenotype. Indeed, here we demonstrated that neutralization of this secreted IL-10 during the activation/expansion of CD8 T cells reversed the increase in specific cytotoxicity and the inhibition of IFN-γ secretion seen in PTPN2KO/PTPN1Het cells, revealing an active role for IL-10 in the acquisition of the phenotype, potentially by mediating combined STAT1 and STAT3 signals. In a recent report, Sun and colleagues found STAT3 to be required for the formation of Tim3+ terminal effectors[43], a highly functional CD8 T cell associated with effective control of infections and tumors, and equivalent to the CX3CR1+ population. Given the reversibility of the phenotype, we believe then that the increase in basal STAT3 phosphorylation comes mainly as result of an autocrine IL-10 loop, instead of lack of regulation by the absence of the phosphatase activity. However, although the signals required for the differentiation of CD8 T cells into IL-10 secreting effectors are not well studied, IL-27 is likely to have a role[83]. IL-27, in turn, also signals through STAT1 and STAT3 in a parallel to IL-10 and IL-24, suggesting that the combined action of PTPN1 and PTPN2 modulate a positive proinflammatory loop involving these cytokines exclusive to CD8 T cells.

Finally, here we demonstrated as also done by others recently [7, 8], that pharmacological inhibition of PTPN1 and PTPN2 is a powerful therapeutic tool when used in combination to immune checkpoint inhibitors, as those targeting PD-1. PTPN1/2 inhibitors offer a safe and cost-efficient non-biological alternative, which activity is not limited to immune cells but also to tumor cells as melanoma[3] and prostate cancer [84].

## Supporting information

All supplementary figures

## Acknowledgments

This work was supported by a Canadian Institute of Health Research Foundation grant (CIHR FDN-159923), the Richard and Edith Strauss Canada Foundation, and the Aclon Foundation to MLT. MLT holds a Chair of the Jeanne and Jean-Louis Levesque Foundation in Cancer Research. LAPQ is the recipient of a Cole Foundation postdoctoral fellowship, ZMC received support from the Quebec provincial MITACS awards, and AP and CHF are recipients of the FRQS training studentships. We thank the “Comparative Medicine and Animal Resources Centre “(CMARC) at McGill University for their veterinarian and animal husbandry services. We are grateful to the Flow Cytometry and Bioinformatics cores at the Rosalind and Morris Goodman Cancer Institute for their help in technical aspects and experimental design. We thank to OncoDesign for the in vivo tumor study with EMT-6 cells.

## Materials and methods

### Mice and cell lines

All mice procedures were performed using 6-12 weeks old C57BL/6 mice according to the Canadian Council on Animal Care ethical regulations and were approved by the McGill University Research and Ethics Animal committee. For conditional deletion in CD8 T cells, mice expressing CRE under the E8i enhancer [46] were bred with mice carrying “floxed” versions of the ptpn1 [45] and ptpn2 alleles [44]. For specific cytotoxicity, mice were crossed with C57BL/6-Tg (TcraTcrb)1100Mjb/J “OT-1” animals purchased from The Jackson Laboratory. For tumor implantation, 6-8 weeks old C57BL/6J male mice were purchased directly from The Jackson Laboratory. Complete necropsies of CRE controls and dKO animals were performed by the Charles River Research Animal Diagnostic Services.

E.G7 cell line, a derivative of the EL4 mouse thymoma cell line expressing chicken ovalbumin (OVA), was purchased from ATCC.

### Antibodies and reagents

For flow cytometry, the following anti-mouse antibodies were acquired from Biolegend CD3 (145-2C11), CD4 (GK1.5), CD8 (53-6.7), TER119 (TER119), Ly-6G/Ly-6C (RB6-8C5), B220 (RA3-6B2), CD11b (M1/70) and KLRG1 (2F1/KLRG1). Antibodies against mouse CD62L (MEL-14) and CD127 (SB/199) were from BD Biosciences. Antibodies against CD44 (1M7) and EOMES (Dan11Mag) were from eBiosciences.

Anti-human antibodies were acquired from BD Biosciences for, CD3 (SK7), CD4 (RPA-T4), CD45RO (UCHL1), CD62L (DREG-56), IFN-γ (4S.B3), TNF-α (MAb11) and ThermoFisher Scientific for CD8 (RPA-T8).

For immunoblotting, the following antibodies were purchased from Cell Signalling Technologies IRF4 (Cat#4948; P173), STAT1 (Cat#9172T), STAT3 (Cat#4904), STAT4 (Cat#2653), STAT5a (Cat#94205), STAT6 (Cat#5397), pSTAT1 Y701 (Cat#7649), pSTAT3 Y705 (Cat#9131), pSTAT4 Y693(Cat#5267), pSTAT5 Y694 (Cat#9351) and pSTAT6 Y641 (Cat#9361). Antibodies against STAT2 (Cat#ab134192) and pSTAT2 Y690 (Cat#ab53132) were from Abcam. The antibody for ShcA was from BD Transduction Laboratories (Cat#S14603). The antibody for BATF3 was from R and D Systems (Cat#AF7437). Antibody against Myb is from Sigma-Aldrich (Cat#05-175, Clone 1-1), Anti-ZEB2 was from Novus Biologicals (Cat#NBP1-82991) and Anti-BLIMP1 was from eBioscience (6D3). Anti-PTPN2 (3E2) was produced in-house [85], Anti-PTPN1 was from Upstate (EMD Millipore, Cat#07-088) and Anti-Calnexin (CNX) was a gift from Dr. John J.M. Bergeron. Kanyr Inc kindly donated PTPN2/PTPN1 specific phosphatase small molecule inhibitors L598 and KQ791.

### Cell purification, activation/expansion and IL-10 neutralization

Cell isolation from splenocytes was performed with StemCell Technologies kits for CD4 (Cat#18952), CD8 (Cat#18953), and CD8 naïve (Cat#18958) following the manufacturer’s instructions. Purified naïve CD8 T cells were seeded on non-tissue culture treated 24 well plates coated with 2mg/ml αCD3ε (Ultra LEAF© purified, clone145-2C11, Biolegend) at 2x10^5^ cells per well, in complete RPMI (RPMI 1640, 10% heat-inactivated FBS, 0.05mM β-mercaptoethanol and 1X Penicillin/Streptomycin Gibco) soluble αCD28 (Ultra LEAF© purified, clone 37.51, Biolegend) and 20U/ml recombinant murine interleukin 2 (rmIL-2) (Peprotech, cat. number 212-12). For stimulation with the cognate OVA 257-264 peptide (SIINFEKL), 2.5 x 10^5^ splenocytes from OT-1 mice were seeded in one ml of complete media in 24 well plates and 0.5µg/ml of peptide (Sigma-Aldrich, cat. number S7951). After 48 hours blasts were expanded with 100 U/ml of rmIL-2 in fresh media and harvested on day four or six after initial stimulation.

In experiments where pharmacological inhibitors were used a 10mM stock solution in PBS was added to the media at the appropriate final concentration. An equal volume of PBS was added to the control cells. The fresh inhibitor was added every 48hs at the corresponding concentrations.

In experiments where IL-10 activity was neutralized, cells were incubated for the period of activation expansion with 20µg/ml of anti IL-10 antibody (JES5-2A5; Biolegend catalog No. 504908) or corresponding isotype control (Biolegend catalog No. 400432).

### Flow cytometry

Spleen, lymph node, and thymus samples were prepared by macerating organs through a 70µm cell strainer (Falcon) to obtain a single-cell suspension; samples were kept on ice in 2%FBS PBS during the staining procedures. For intracellular staining of nuclear factors, samples were processed with the FOXP3/Transcription Factor Staining Buffer Set (eBioscience, Thermo-Fisher) following the manufacturer’s instructions. Samples were read in a BD LSR Fortessa instrument and analyzed with FlowJo software.

### Human T cell isolation and invitro expansion

Peripheral blood mononuclear cells (PBMCs) of healthy donors were obtained after informed consent approved by the local ethics committee (Hôpital Maisonneuve-Rosemont, Montreal, Québec; protocol #2020-2141) by venipuncture followed by manual (Ficoll-Paque, GE Healthcare, Canada) gradient density separation. T cells are isolated using an EasySep™ Human T Cell Enrichment Kit as described by the manufacturer (Stemcell Technologies). For T cell activation, 96 well plates were coated with anti-CD3 (Clone #145-2C11, BD Biosciences) antibody at five µg/mL for 1h30 at 37C. After washing with PBS, 100,000 PBMC per well were seeded in complete media (Advanced RPMI with 10% human serum and 1% L-Glutamine) supplemented with 1µg/mL of anti-CD28 antibody (Clone #37.51, BD Biosciences). The L598 compound was added to the cultures every two days at 2, 5, 10, or 25µM. After 7 days, cells are harvested for analysis. Phenotypic evaluation of T cells is performed to determine the differentiation status. For functional analysis, T cells were incubated in 96 well plates with Brefeldin A (15µg/mL, Biolegend) and stimulated with PMA (5ng/mL, Sigma) and Ionomycin (500ng/mL, Sigma) for 4 hours at 37C. Cells were harvested, stained for extracellular markers, and then permeabilized (eBioscience™ Foxp3 / Transcription Factor Staining Buffer Set) overnight at 4C. After permeabilization, cells were washed and stained with TNF-α and IFN-γ antibodies to determine cytokine production following stimulation. Flow cytometry acquisition was performed on the BD LSRII instrument.

### In vitro CTL-specific cytotoxicity

For specific cell cytotoxicity, a flow cytometry-based assay was used. EG.7 target cells were stimulated 24 hours before the experiment with 100 U/ml of rmIFN-γ to increase the expression of Class I molecules. OT-1 CD8 T cells were activated/expanded as mentioned above and harvested on day 4 for experiments. In experiments involving pharmacological inhibitors, cells were washed twice with media to prevent any side effects on target cells. EG.7 target cells were loaded with CellTracker™ Orange CMRA dye (ThermoFisher Scientific) for 30 min, following the manufacturer’s instructions. After being washed extensively with serum-containing media to inactivate the dye remaining in the solution. After, CD8 effector and EG.7 target cells were plated at the corresponding ratios in round bottom 96 well plates, centrifuged at 300g for 2 min, and incubated a 37°C and 5% CO2 for 5 hours. Some wells were plated with target cells alone to be used as spontaneous dead controls. Following incubation, cocultures were centrifuged for 5 minutes at 300g, and the supernatant was harvested for cytokine quantification. Cocultures were then washed with PBS and stained with LIVE/DEAD™ Fixable Blue Dead Cell Stain Kit or LIVE/DEAD™ Fixable Aqua Dead Cell Stain Kit (ThermoFisher Scientific) following the manufacturer’s instructions. Cocultures were then incubated for 30 minutes protected from light, and washed twice with 2%FBS PBS. After the last wash cocultures were either read immediately or fixed with 2% PFA in PBS for 45 min, washed, and resuspended in 2%FBS PBS.

Samples were acquired in BD LSR Fortessa or BD Canto machines and analyzed with FlowJo software. The target cell death percentage was calculated as the percentage of LIVE/DEAD stain-positive cells from the CMRA-positive events. Spontaneous cell death was subtracted from that observed in experimental samples to calculate specific target cell cytotoxicity. Values correspond to the average of technical replicates. This protocol was optimized to obtain a non-saturated curve from the samples with the highest activity (N2KO/N1Het and dKO CTLs), hence the lower values obtained with CRE controls.

### CTL stimulation with INF-β, protein extraction, and immunoblotting

To evaluate protein phosphorylation in CD8 T cells after stimulation with IFN-β (Peprotech), day 4 CTL were washed twice in serum-free media and incubated for 2 hours under serum and cytokine deprivation before stimulation; serum-free media was maintained during stimuli. To stop stimulation cells were lysed by adding an equal volume of ice-cold 2X RIPA buffer (100 mm Tris– HCl pH 7.4, 200 mm NaCl, 0.2 mm EDTA, 2% NP40, 0.2% SDS) containing protease (Roche complete protease inhibitor cocktail, Sigma-Aldrich) and phosphatase inhibitors (4mM sodium orthovanadate, 10mM NaF, 20 mM sodium pyrophosphate). Lysates migrated through 10 or 12% PAGE and transferred to PVDF membranes (Immobilon-P, EMD Millipore).

### ELISA

To measure cytokine secretion in supernatants from cytotoxicity experiments, enzyme-linked immunoassays (ELISA) for mouse IFN-γ Biolegend (Cat#430804) and IL-10 ThermoFisher (Cat#88-7105-88) were used according to manufacturer’s instructions.

### mRNA isolation

To reduce potential variations given by the differentiation state of splenic CD8 T cells, mRNA for bulk sequencing experiments was isolated from day 4 CTLs expanded from purified splenic naïve CD8 T cells. For each sample, 2x 10^6^ cells were processed under laminar flow cabinets with the RNAEasy plus mini from Qiagen (Cat# 74134) following the manufacturer’s instructions.

### Gene expression profiling

After RNA extraction, libraries for sequencing were prepared using the Illumina TruSeq RNA Library Prep Kit following the manufacturer’s instructions. High throughput sequencing was performed at the Centre d’expertise et de services Génome Québec. 28 to 46 million reads were obtained per sample, averaging 35 million reads.

### mRNA sequencing data processing

The resulting reads were aligned to the GRCm38 mouse reference genome assembly, using STAR [86]. SAM files were sorted with samtools 1.10. Read counts were obtained using HTSeq [87] with parameters -m intersection-nonempty -stranded=reverse.

For all downstream analyses, we excluded lowly-expressed genes with an average read count lower than ten across all samples. Raw counts were normalized using edgeR’s TMM algorithm [88] and were then transformed to log2-counts per million (log CPM) using the voom function implemented in the limma R package [89]. We fitted a linear model using limma’s lmfit function to assess differences in gene expression levels. Nominal p-values were corrected for multiple testing using the Benjamini-Hochberg method. Gene set enrichment analyses were performed using Enrichr [90]. Volcano/dot plots and heatmaps were created using the R package ggplot2 and Complex Heatmap, respectively.

### Invitro phosphatase assay

Enzyme reactions were conducted in assay buffer 50mM HEPES pH7.0, 3mM DTT and 1mg/mL BSA using the purified recombinant GST-PTP1B and GST-TC-PTP catalytic domains [91]. DiFMUP was used as a substrate.

Determination of kinetics constants using DiFMUP as substrate: The hydrolysis of DiFMUP was conducted in black 96-well plates (Corning) in a final volume of 100µL at 25°C. The reaction was monitored by measuring the fluorescence (excitation wavelength 358nm/emission 455nm) with the Varioskan plate reader (Thermo Electron). Enzyme dilution was determined by choosing a reaction rate to comprise in an Absorbance range of 5-30 FU units/min. For the kinetic assays fluorescence was monitored over 10 minutes in 30 seconds intervals and rates were calculated using a non-linear least-square fitting procedure. Km was determined from rates at various substrate concentrations using the Michaelis-Menten equation and IC50 values were derived by a sigmoidal dose-response (variable slope) curve using GraphPad Prism software. A substrate concentration equivalent to the Km value was used for IC50 determinations. KQ-791 and L598 Inhibitors were diluted in PBS, and then a serial dilution starting from 24µM and covering 3 log scales was made in assay buffer.

### Tumor inoculation and immunotherapy models

For immunotherapy models, 8-10 weeks old BALB/c females were purchased from The Jackson laboratory. Mice were shaved in the right flank and 10^6^ EMT-6 cells were injected in 100 µl of PBS. Tumor volume was calculated by measuring 2 axes (length and width) and applying the formula V = 4/3π × [(Length/2) × (Width/2)2]. For treatment intraperitoneal injections of KQ-791 at 250µg/g of weight and/or 10µg/g of weight of anti-PD-1 antibody, Bio X Plus (Clone RPM1-14; Cat# BP0146) or Isotype Control were started at day 11 post tumor inoculation after randomization. KQ-791 (kind gift from Kanyr Pharma Inc.) injections of KQ791 were repeated every 2 days for a total of 10 doses, and anti-PD-1 or isotype control injections were repeated twice every week for a total of 3 weeks. Humane intervention for euthanasia followed McGill University Research and Ethics Animal committee guidelines, including tumor volume reaching 2000 mm3, tumor ulceration or deterioration of the clinical condition. Only mice reaching humane endpoint by tumor volume were included in the experiments.

### Statistical analysis

All statistical analyses were performed using GraphPad Prism software.

## Notes

https://www.ncbi.nlm.nih.gov/bioproject/PRJNA869093

### Competing Interest Statement

Michel L. Tremblay acts as Chief Scientific Officer and Luis Alberto Perez-Quintero acts as Research Project Director for Kanyr Pharma inc. Both MLT and LAPQ have a current patent on the use of KQ791 for systemic immunotherapy. No other authors declare any conflict of interest.

### Summary of Updates

Major revision of the introduction and main body, new data on the role of IL-10 in the phenotype acquired, and the synergy of combination therapy with PTPN1/2 inhibitors and with anti-PD1 immune blockade agents in breast cancer model. Small revision of other sections and formatting to comply with JCI standards.

