## Supplementary material for "PTPN1/2 inhibition induces highly functional terminal effector CD8 T cells through autocrine IL-10": All supplementary figures

A

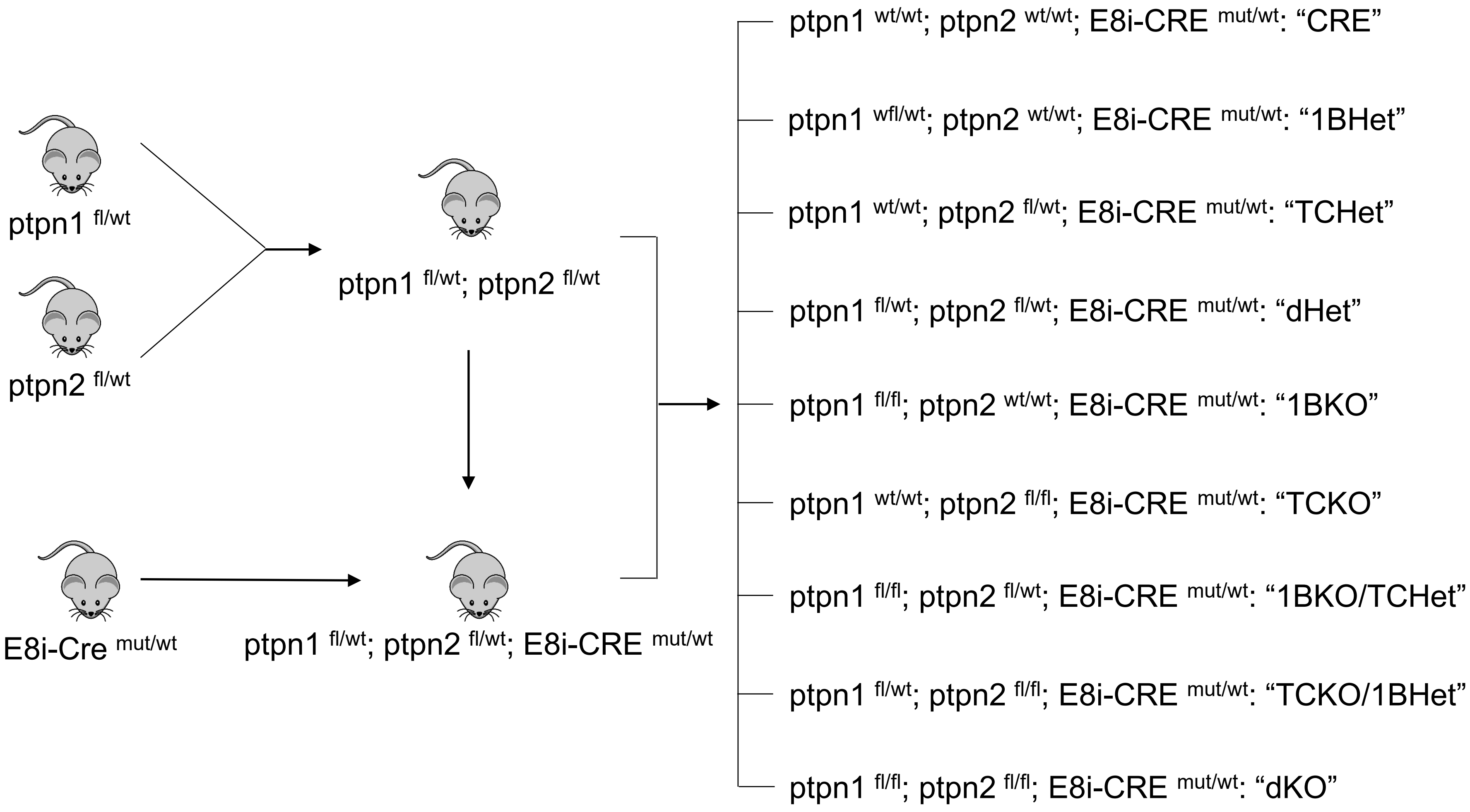


Figure S1

(Continue next page)

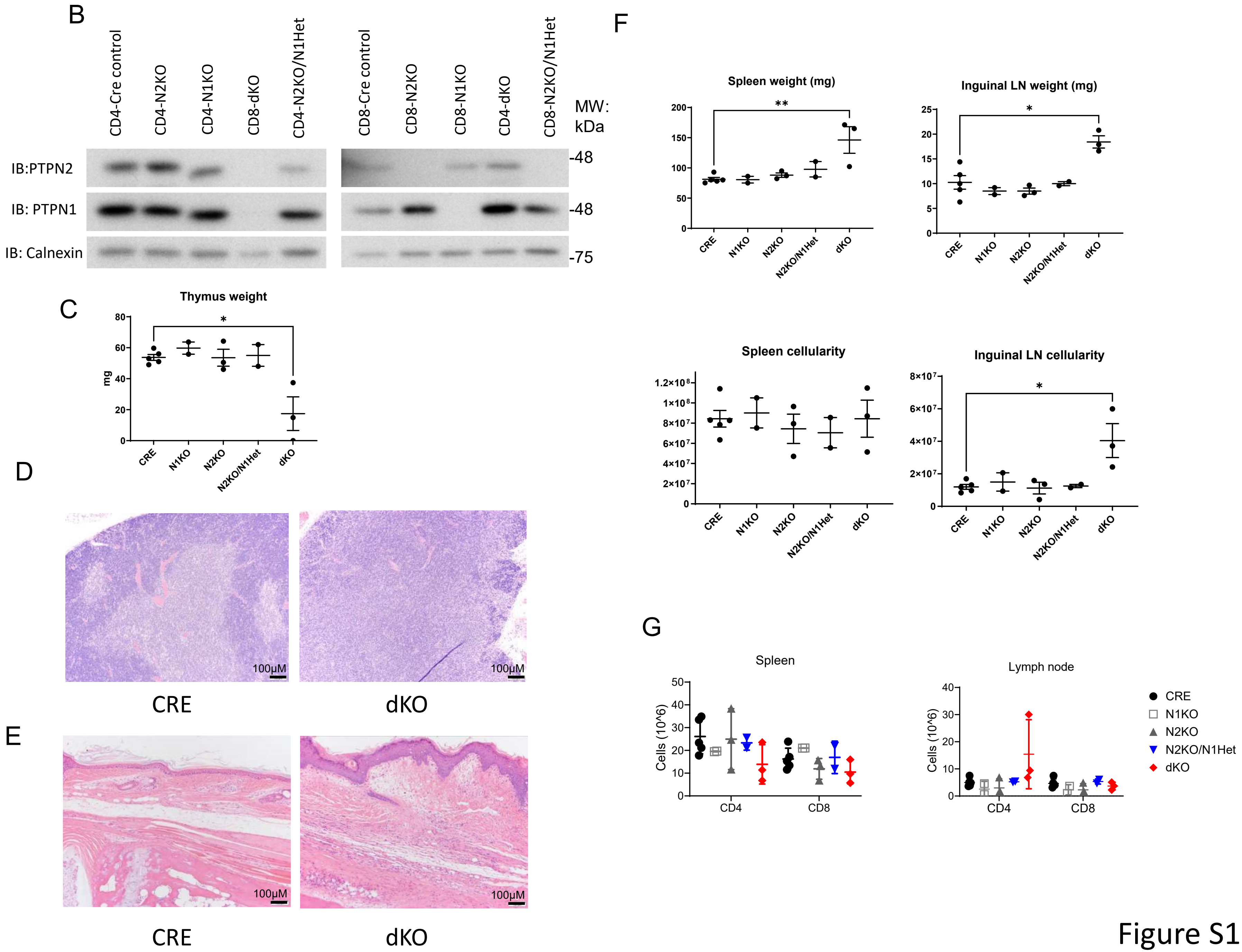

B
CD4-Cre control
CD4-N2KO
CD4-N1KO
CD8-dKO
CD4-N2KO/N1Het
CD8-Cre control
CD8-N2KO
CD8-N1KO
CD4-dKO
CD8-N2KO/N1Het
MW: kDa
IB:PTPN2
IB: PTPN1
IB: Calnexin
-48
-48
-75
C
Thymus weight
mg
CRE
N1KO
N2KO
N2KO/N1Het
dKO
D
100μM
CRE
dKO
E
100μM
CRE
dKO
F
Spleen weight (mg)
Inguinal LN weight (mg)
Spleen cellularity
Inguinal LN cellularity
G
Spleen
Lymph node
Cells (10^6)
CD4
CD8
CRE
N1KO
N2KO
N2KO/N1Het
dKO


Figure S1

**Figure S1. Conditional deletion of TC-PTP and PTP1B in CD8 T cells.** A) Breeding schematics showing the steps to obtain the different genotypes analysed. B) Immunoblot for TC-PTP, PTP1B and Calnexin as loading control from protein homogenates of CD4 and CD8 T cells with different genotypes analysed. C) Thymus size, expressed as weight, from 8–12-week-old male mice carrying the different genotypes analysed. D) and E) Hematoxylin and eosin staining of D) thymus and E) skin sections from 10-week-old CRE controls or TC-PTP/PTP1B (dKO) CD8 specific "knockouts". F) Spleen and inguinal lymph node size (by weight) and cellularity from the different genotypes analysed. G) Total number of CD4 and CD8 T cells from spleen and inguinal lymph nodes shown in Figure 1B, no significant differences were found. (Bars represent +/- S.E.M.; One-way (C, E) or Two-way (F) ANOVA; P values *<0.05, **<0.005, ***<0.0005, ****<0.00005).

Figure S1

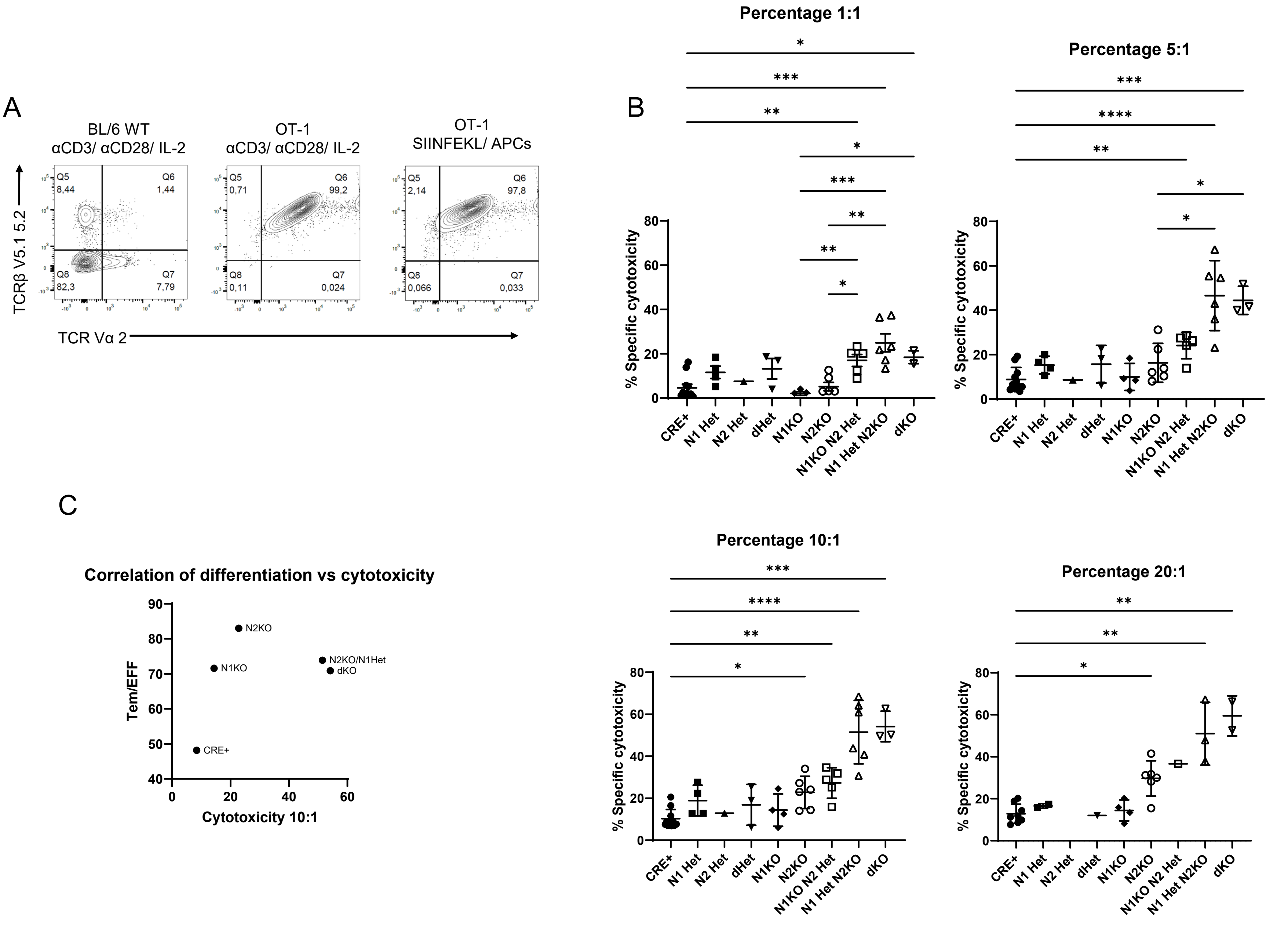
A
BL/6 WT
αCD3/ αCD28/ IL-2
OT-1
αCD3/ αCD28/ IL-2
OT-1
SIINFEKL/ APCs
TCRβ V5.1 5.2
TCR Vα 2
Q5 8,44
Q6 1,44
Q8 82,3
Q7 7,79
Q5 0,71
Q6 99,2
Q8 0,11
Q7 0,024
Q5 2,14
Q6 97,8
Q8 0,066
Q7 0,033
B
Percentage 1:1
Percentage 5:1
Percentage 10:1
Percentage 20:1
% Specific cytotoxicity
CRE+
N1 Het
N2 Het
dHet
N1KO
N2KO
N1KO N2 Het
N1 Het N2KO
dKO
C
Correlation of differentiation vs cytotoxicity
Tem/EFF
Cytotoxicity 10:1
N2KO
N1KO
N2KO/N1Het
dKO
CRE+

Figure S2

**Figure S2. In vitro cytotoxicity against specific target cells of CD8 T cells with genetic deficiency of PTPN2 and/or PTPN1.** A) Purified naïve CD8 T cells from C57Bl/6 WT or OT-1 mice were stimulated to generate activated/expanded CD8 T cells either by receptor crosslinking of CD3 and CD28 with specific antibodies and IL-2 (leftmost and central plots) or with cognate peptide and APC (right plot). Cells were analyzed by flow cytometry for the expression of the TCR variable alpha chain two and TCR beta variable chain 5.1. Cells expressing the transgenic chains in the OT-1 mouse are positive for both (Q6). B) Statistical representation of day 4 activated/expanded CD8 T cells from all the different genotypes obtained, tested for specific cytotoxicity against the OVA expressing E.G7 thymoma cells. Each panel represents a different effector to target ratio. To simplify statistical differences, panels for 5:1, 10:1, and 20: ratios show comparisons of all groups to CRE and N2KO groups (Bars represent +/- S.E.M.; Kruskal-Wallis test; P values *<0.05, **<0.005, ***<0.0005, ****<0.00005) C) Correlation plot of effector memory/effector differentiation against cytotoxic activity against E.G7 cells at 10 to 1 ratio for the different genotypes analyzed. Spearman's rank correlation was computed to assess the relationship between both variables, and no correlation was observed (r=0.3, p=0.3417).

Figure S2

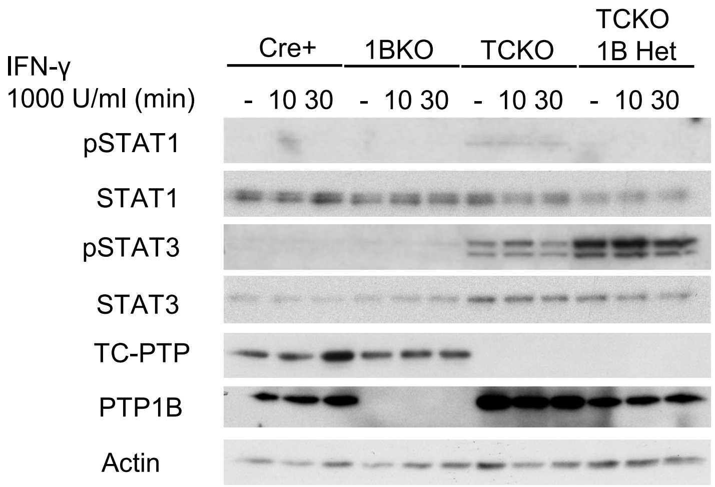


**Figure S3. In vitro stimulated/expanded CD8 T cells are unsensitive to INF-γ.** Immunoblotting of protein lysates from day 4 post activation/expansion CD8 T cells, stimulated with 1000 Units of recombinant murine interferon γ (rmIFN-γ) for the indicated times. Unstimulated controls (-) correspond to cells without the addition of the cytokine. Membranes were reprobed for Actin as loading control (Representative of at least 3 independent experiments).

Figure S3

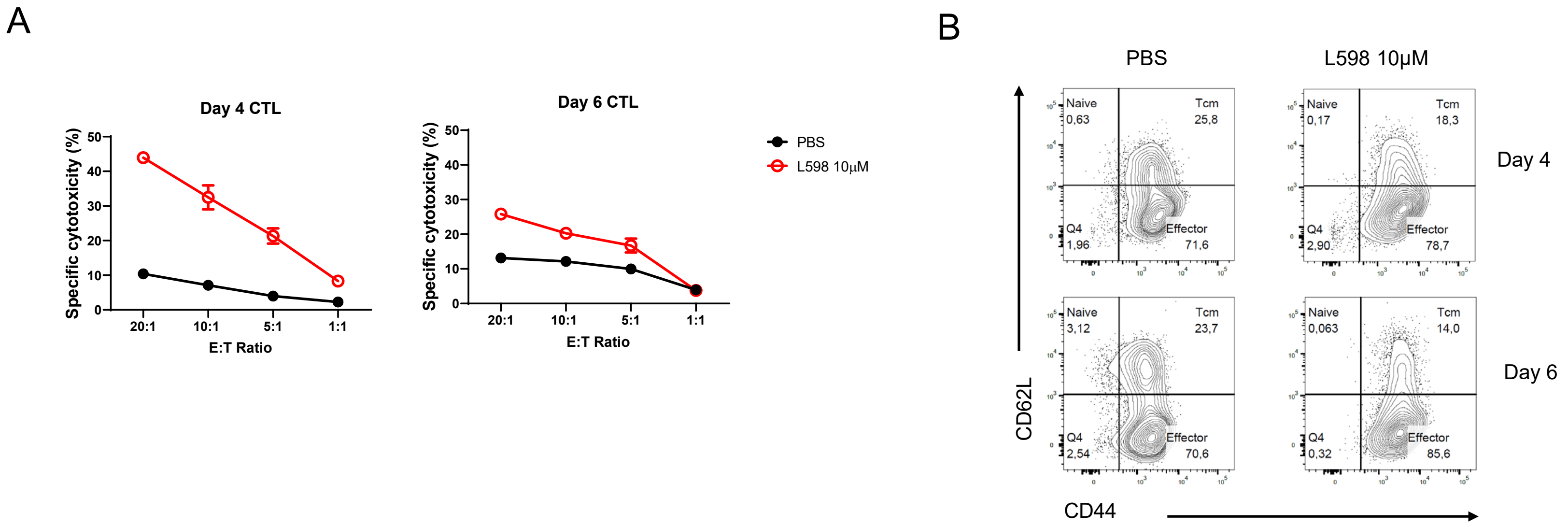


**Figure S4. L598 treatment enhances effector differentiation in mouse T cells.** Splenocytes from OT-1 mice were stimulated with the OVA peptide 257-264 (SIINFEKL) for two days and later expanded with IL-2, in the presence of 10μM L598 or the vehicle (PBS) as control. A) Day 4 and day 6 CTL were tested for specific cytotoxicity against E.G7 OVA-expressing thymoma cells. B) Cells in A were assessed for expression of the differentiation markers CD62L and CD44. (Representative of 2 different experiments).

Figure S4

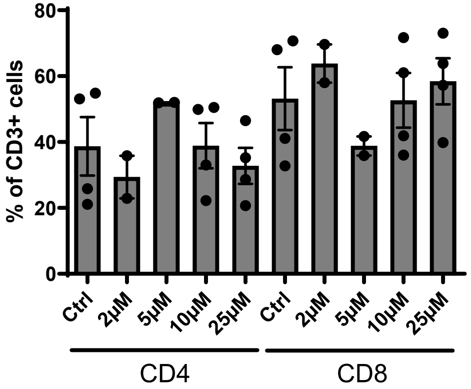


**Figure S5. L598 treatment does not affect the CD4/CD8 ratio of human T cells.** Purified T cells from human donors were activated by αCD3/αCD28. Percentual contribution of CD4 and CD8 T cells of in vitro human T cell cultures at day 7.

Figure S5

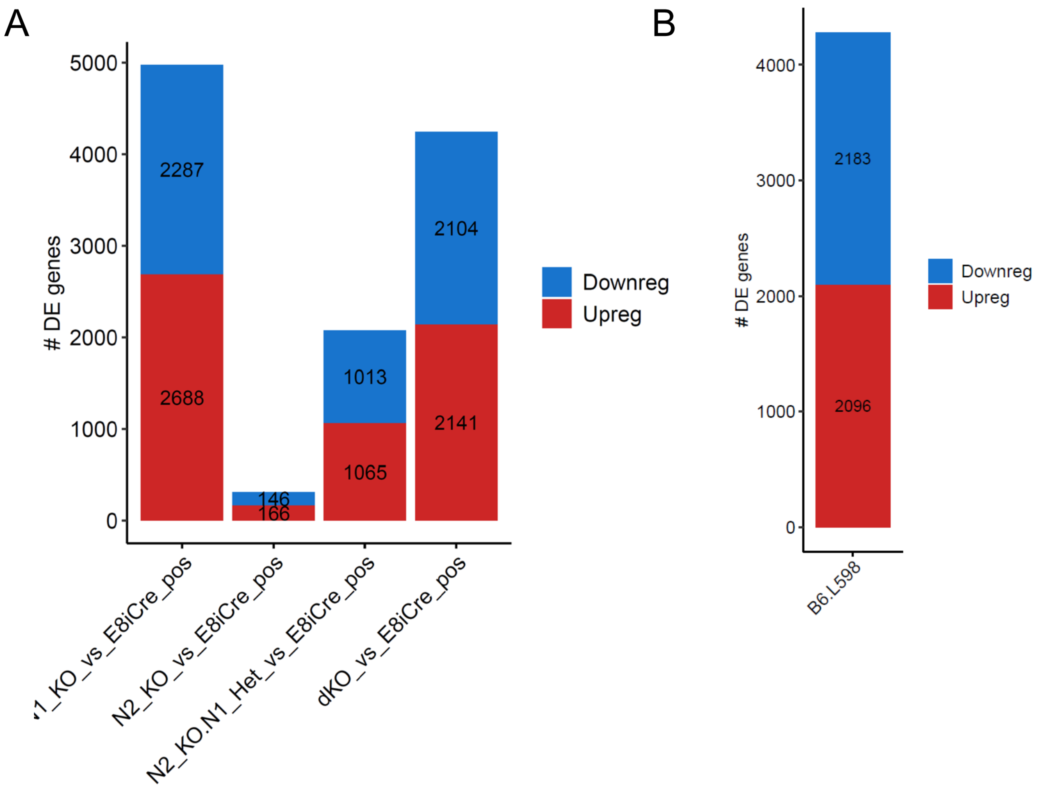


**Figure S6. Bulk mRNA analysis of CD8 T cells with reduced activity of PTPN1 and PTPN2.** A) Number of differentially expressed (DE) genes in day 4 activated/expanded CD8 T cells compared to CRE controls. B) Number of DE genes in in day 4 activated/expanded CD8 T cells from L598 treated cells compared to controls.

Figure S6

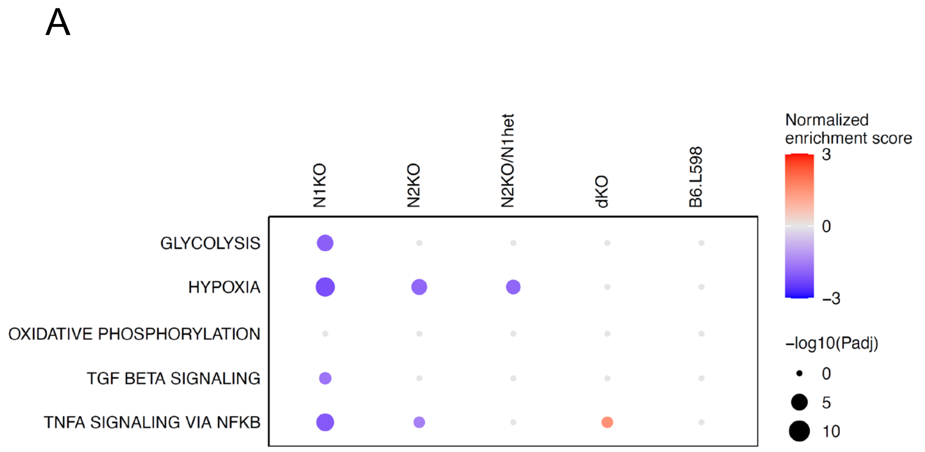


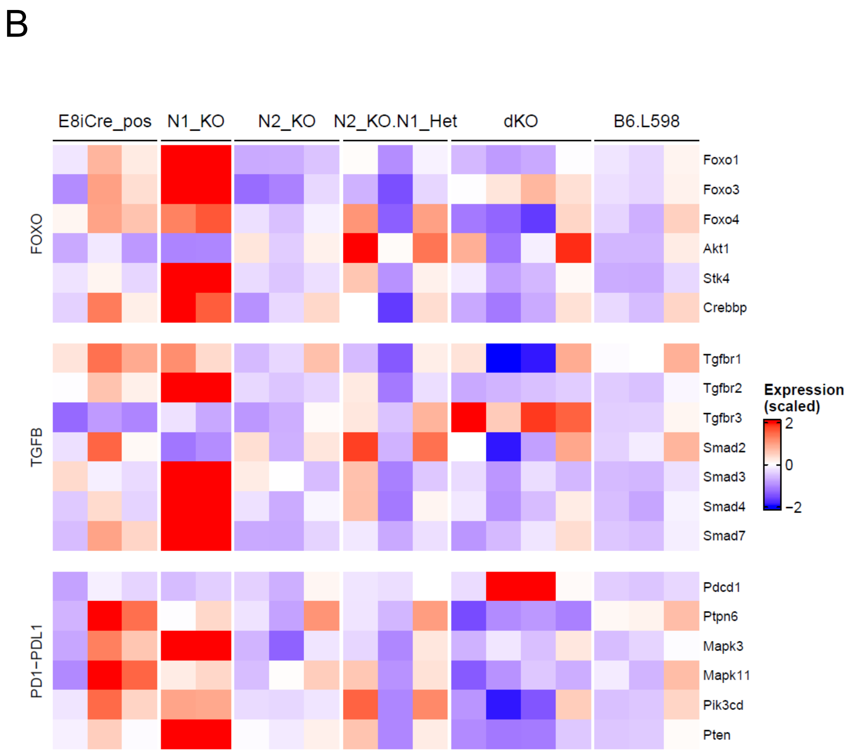


**Figure S7.** A) List of additional significantly scoring Hallmark collection gene sets not shown in Figure 6. B) Heat map of individual genes scores given by pathfinder analysis for the FOXO, TGF-β and PD1-PDL1 pathways.

A

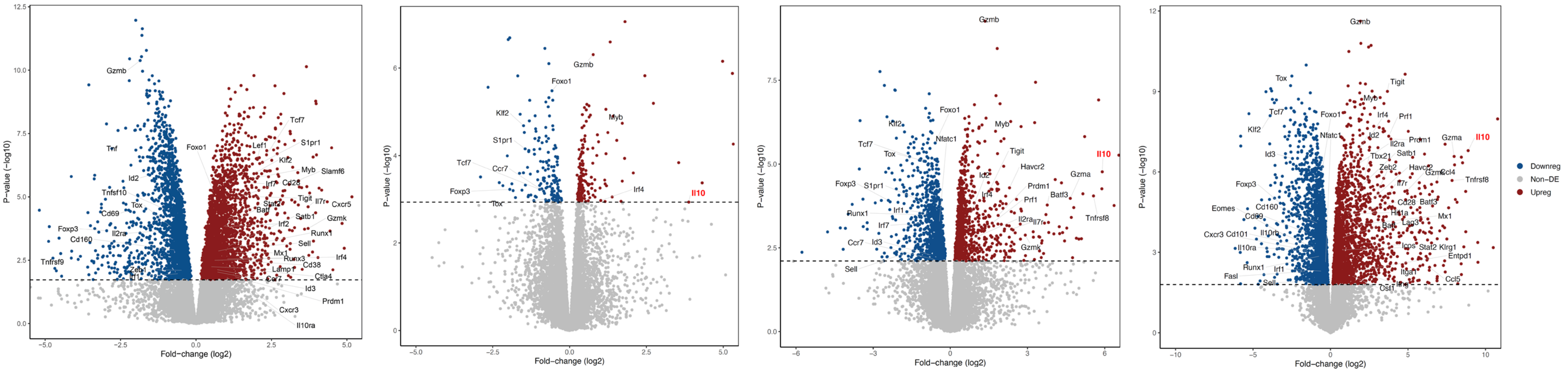


N1KO vs. CRE+

N2KO vs. CRE+

N2KO/N1Het vs. CRE+

dKO vs. CRE+

B

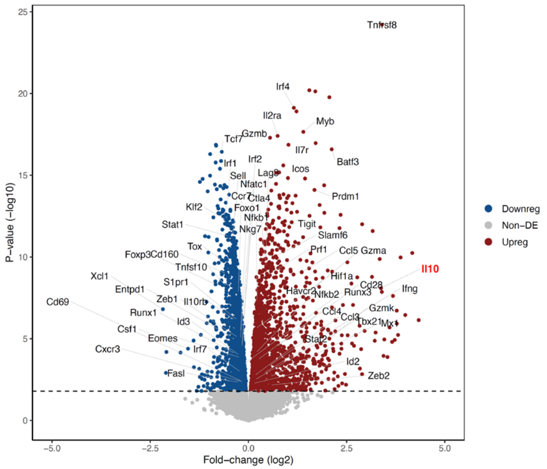


L598 vs. B6

**Figure S8. IL-10 transcript is upregulated upon PTPN1 and PTPN2 inhibition.** Volcano plots representing the DE genes of A) the significant differentially expressed genes between day 5 CTLs from CRE controls and the different genotypes analyzed and B) the significant differentially expressed genes between day 5 CTLs from C57BL/6 mice, treated or not with 5µM L598. Red squares highlight the expression of the IL-10 transcript.

Figure S8
