## Supplementary table 1 for "PTPN1/2 inhibition induces highly functional terminal effector CD8 T cells through autocrine IL-10"

| Upregulated | | |
|---|---|---|
| Gene | log2FC | -LOG10 Pv |
| Gm16867 | 4.583613765 | 7.743969558 |
| Arhgef10l | 3.375523703 | 3.162380839 |
| Anxa3 | 3.323887376 | 4.370440005 |
| Trp73 | 3.183864169 | 4.329619822 |
| Il13ra1 | 3.148069558 | 3.347919405 |
| Sval1 | 3.113378248 | 6.97800749 |
| Maats1 | 3.073868754 | 3.509705509 |
| Hlx | 3.017016953 | 11.62988197 |
| Fabp4 | 2.990486107 | 5.730548812 |
| Gad2 | 2.884878671 | 3.033409812 |
| Gm33994 | 2.85130388 | 3.282265842 |
| Akap6 | 2.805937074 | 4.076282813 |
| Gm17590 | 2.677061312 | 6.880135096 |
| Gm13546 | 2.511571834 | 10.62700473 |
| Moxd1 | 2.505851231 | 4.898137715 |
| Arhgap6 | 2.475686655 | 3.287223596 |
| Gpx8 | 2.461602554 | 3.651273782 |
| Gm34342 | 2.421787499 | 3.649760992 |
| Mycl | 2.42133903 | 3.496987024 |
| Bcam | 2.401218391 | 2.876069905 |
| Cox6b2 | 2.358188136 | 9.28164924 |
| Plcd1 | 2.338691483 | 4.580231142 |
| Bmerb1 | 2.325565928 | 5.090164095 |
| Tnfrsf8 | 2.21959723 | 8.143064317 |
| Ahrr | 2.171479872 | 2.958603479 |
| Il10 | 2.065970066 | 4.763881612 |
| Kif1a | 1.978605223 | 2.893025761 |
| Pparg | 1.931965146 | 3.015266143 |
| Cpd | 1.902139606 | 6.356393186 |
| Cass4 | 1.896038073 | 3.027369987 |
| Ifitm7 | 1.808039402 | 5.578813721 |
| Penk | 1.767763324 | 4.047145185 |
| Padi4 | 1.649125986 | 5.329849139 |
| Batf3 | 1.630569268 | 4.543886131 |
| Gm5111 | 1.529710663 | 2.835285989 |
| Fndc3c1 | 1.501048629 | 3.070623856 |
| Map3k20 | 1.486652588 | 3.231948298 |
| Il7r | 1.465371972 | 4.335839407 |
| Slc24a3 | 1.41855724 | 4.047935999 |
| Nectin2 | 1.401208294 | 3.446159076 |
| Mt3 | 1.40120788 | 3.305588471 |

| Downregulated | | |
|---|---|---|
| Gene | log2FC | -LOG10 Pv |
| Gas2l3 | -0.502012194 | 2.980671302 |
| Acss1 | -0.506466047 | 4.573504106 |
| Nktr | -0.507405419 | 5.347222359 |
| Bicra | -0.510591592 | 5.445172534 |
| Hemgn | -0.51127167 | 4.547699236 |
| Tle4 | -0.520268744 | 6.7620124 |
| Sesn1 | -0.524984258 | 4.012194644 |
| Prex1 | -0.528346061 | 6.951378128 |
| Ccdc50 | -0.537479406 | 3.905213962 |
| Ash1l | -0.540294946 | 3.080580855 |
| Tnfrsf26 | -0.542198363 | 3.33295241 |
| 2310043P16Rik | -0.549594177 | 4.31760093 |
| Gata3 | -0.558759021 | 2.969691432 |
| Tcf7 | -0.565319484 | 3.607971546 |
| 2610005L07Rik | -0.566123816 | 3.339881344 |
| Evl | -0.583454136 | 5.369978764 |
| Gm3200 | -0.593380172 | 2.87093051 |
| Dync1h1 | -0.597933278 | 3.585904025 |
| Cd27 | -0.600331588 | 3.486227603 |
| Skp2 | -0.604852641 | 3.720076116 |
| Ehd3 | -0.608487427 | 3.245810206 |
| Fam117a | -0.609620031 | 3.65232734 |
| Tmem154 | -0.626271396 | 3.228257557 |
| Lca5 | -0.629018682 | 3.0906724 |
| Elovl7 | -0.635050232 | 4.578446986 |
| Tbxa2r | -0.637309936 | 2.931944483 |
| G2e3 | -0.655659246 | 6.102781899 |
| Edaradd | -0.656367868 | 3.607642471 |
| Zfp712 | -0.657208152 | 3.450530626 |
| Zfp831 | -0.661319463 | 3.655008638 |
| Ripor2 | -0.673148863 | 3.669836684 |
| Plxdc2 | -0.696129569 | 5.680476542 |
| Lrig1 | -0.699182853 | 3.096745954 |
| Sfmbt2 | -0.709083502 | 2.969245871 |
| C330011M18Rik | -0.721230998 | 3.352165289 |
| 2610035D17Rik | -0.723291251 | 3.015218775 |
| Vps37b | -0.724687709 | 3.751745276 |
| Prr5l | -0.726477405 | 3.056822748 |
| Klf2 | -0.745458141 | 3.218615173 |
| Tmem71 | -0.77367078 | 3.19317443 |
| Pygm | -0.79206733 | 4.074225861 |

| | | |
|---|---|---|
| Scn1b | 1.383081994 | 2.899839631 |
| Dab2ip | 1.381460458 | 3.578651187 |
| B4galnt4 | 1.374246782 | 3.765898243 |
| Gorasp1 | 1.267787994 | 2.945807965 |
| Src | 1.243540336 | 3.651586607 |
| Esm1 | 1.223895897 | 4.414316024 |
| Galr3 | 1.216936209 | 3.483769225 |
| Fcmr | 1.182039921 | 3.261592659 |
| Naprt | 1.157798438 | 3.039930874 |
| Prf1 | 1.1519856 | 6.040342594 |
| Cdkn1a | 1.151980185 | 5.561611169 |
| B3gnt8 | 1.121069036 | 3.005041539 |
| Cytip | 1.105051007 | 6.75887093 |
| Eng | 1.092319552 | 3.247219424 |
| Fam234a | 1.089557153 | 6.950769694 |
| Ube2e2 | 1.083067254 | 3.036032539 |
| Tgfb3 | 1.0802005 | 5.152952718 |
| Mansc1 | 1.046624422 | 2.958537624 |
| Cacnb3 | 1.028047022 | 3.757447742 |
| P2ry2 | 1.025250499 | 6.207680359 |
| Slc4a8 | 1.021291468 | 4.901066972 |
| Syngr3 | 1.020813681 | 3.03413376 |
| 9330159M07Rik | 0.989205708 | 2.961141912 |
| Spsb1 | 0.977316922 | 3.43816766 |
| Adcy6 | 0.975765765 | 3.538104513 |
| Serpine2 | 0.9595418 | 5.675891263 |
| Coprs | 0.948884515 | 3.983246787 |
| Nkain1 | 0.942858289 | 3.143284782 |
| Bcl3 | 0.935024701 | 7.346224882 |
| Fosl1 | 0.931285826 | 3.164408109 |
| Ncf2 | 0.918292719 | 4.454399912 |
| Bmpr1a | 0.901075792 | 3.004481144 |
| Trim45 | 0.890603315 | 3.599753354 |
| Slc7a7 | 0.874414152 | 3.287726123 |
| St3gal1 | 0.869173087 | 3.915425675 |
| Mt2 | 0.868944283 | 7.387405918 |
| Gm10602 | 0.862021765 | 2.901466211 |
| Rps13-ps1 | 0.854341549 | 5.079922059 |
| Extl1 | 0.84164927 | 3.402492628 |
| Cerox1 | 0.840221668 | 2.860174757 |
| Cetn4 | 0.809946418 | 5.835656032 |
| Lhx2 | 0.806602262 | 2.841804245 |
| Hoxb4 | 0.796344093 | 3.127512929 |

| | | |
|---|---|---|
| Ticam2 | -0.797342526 | 3.006012632 |
| Bend5 | -0.814435401 | 2.930843817 |
| Il10ra | -0.818019273 | 2.90102625 |
| Aff3 | -0.81977817 | 5.856725177 |
| Gm2682 | -0.826530698 | 4.637658597 |
| Slc28a2b | -0.830006167 | 3.483473322 |
| Rasa3 | -0.834813988 | 2.946288421 |
| Pisd-ps2 | -0.839848257 | 3.582906453 |
| Npas2 | -0.839851545 | 2.867165018 |
| Fam49a | -0.843794594 | 2.85893436 |
| Gbp7 | -0.84533622 | 2.821395256 |
| 4732416N19Rik | -0.887838274 | 3.307512812 |
| Nsg2 | -0.896069574 | 3.297505974 |
| Samd9l | -0.93548351 | 4.449960914 |
| Mturn | -0.975223514 | 3.170708403 |
| Pdcd4 | -0.986973175 | 5.039986083 |
| 3110083C13Rik | -0.995584885 | 3.148464584 |
| Ighj1 | -0.996132838 | 2.890041807 |
| Serinc3 | -1.009329127 | 10.08134391 |
| Ms4a4c | -1.012774697 | 4.095264999 |
| Gm16124 | -1.038326882 | 3.420714356 |
| Atp1b1 | -1.04206507 | 3.358770758 |
| Cnga1 | -1.066866358 | 3.636991702 |
| Gm38077 | -1.085446479 | 3.270584078 |
| Itih5 | -1.112824792 | 4.344746484 |
| Txk | -1.128772378 | 3.459014607 |
| Ccdc62 | -1.17458307 | 2.815343292 |
| Gm49553 | -1.185698343 | 2.891114556 |
| Gm6093 | -1.203669136 | 4.774717675 |
| Acoxl | -1.225683107 | 3.467275208 |
| Cbfa2t3 | -1.25674333 | 4.298013863 |
| Gm10175 | -1.263640206 | 2.8972074 |
| Gamt | -1.281189299 | 3.04646203 |
| Gm44164 | -1.318519045 | 4.019208273 |
| Eomes | -1.343110395 | 3.208578196 |
| Nipal1 | -1.344039075 | 4.02883483 |
| Sell | -1.542214664 | 5.457026818 |
| A930004J17Rik | -1.609553763 | 3.879302468 |
| Gm44667 | -1.632264275 | 4.970509199 |
| Anp32-ps | -1.65845763 | 5.920730574 |
| Zfp395 | -1.698482198 | 4.460815888 |
| Cd22 | -1.769992673 | 3.473398644 |
| Gm43039 | -1.819054825 | 3.550607112 |

| | | |
|---|---|---|
| Cdkl2 | 0.786086703 | 3.010734896 |
| Abi3 | 0.783150682 | 3.191663963 |
| Rps15a-ps8 | 0.77846786 | 3.04598742 |
| Homer3 | 0.771121613 | 2.904227949 |
| Havcr2 | 0.769882296 | 3.507481975 |
| Phlda3 | 0.764764998 | 3.360752363 |
| Psrc1 | 0.750599027 | 6.242375241 |
| Bbs7 | 0.74941585 | 3.478270351 |
| Chst11 | 0.748546178 | 4.686677155 |
| Dmwd | 0.747073464 | 3.52501326 |
| Echdc2 | 0.745425102 | 3.635691573 |
| Skap2 | 0.737440531 | 5.105850671 |
| Casp3 | 0.735827676 | 6.143266592 |
| Tmem159 | 0.729690281 | 4.339219791 |
| Lilr4b | 0.717937486 | 2.948727871 |
| Zfp365 | 0.699887991 | 3.643817864 |
| Gm14584 | 0.697536502 | 2.992010646 |
| Zc2hc1a | 0.696630744 | 3.260631195 |
| Eps8 | 0.686696773 | 3.03541973 |
| Casp4 | 0.67784227 | 2.803458909 |
| Rgs12 | 0.673342739 | 5.560665089 |
| 2210408F21Rik | 0.6622772 | 4.768845917 |
| Smim3 | 0.661841995 | 4.356889859 |
| 6330418K02Rik | 0.659476606 | 2.976640079 |
| Pmepa1 | 0.630091224 | 2.807077106 |
| Zbtb7b | 0.624941416 | 3.546211461 |
| Arhgef12 | 0.598983226 | 2.897879644 |
| Tnfrsf13b | 0.597515641 | 3.799701922 |
| Edem2 | 0.593816347 | 6.471693718 |
| Zmat3 | 0.579781 | 3.573655205 |
| Gzmb | 0.579191902 | 8.712642968 |
| Il2ra | 0.559423237 | 3.493203021 |
| Fam210b | 0.552334208 | 4.349319118 |
| Zfp944 | 0.551311004 | 2.932678848 |
| Klrk1 | 0.545885162 | 4.211043679 |
| Inka2 | 0.545082195 | 3.050685196 |
| Gngt2 | 0.535558689 | 2.832004099 |
| Aopep | 0.524521532 | 4.92081886 |
| Alcam | 0.502525084 | 4.119845954 |
| Kcnd1 | -1.930844881 | 4.383333037 |
| Rps13-ps2 | -1.964114273 | 4.537311593 |
| Rasgef1a | -1.986251633 | 3.521402607 |
| Gm43300 | -1.995354877 | 4.75838721 |
| Trim34a | -2.012518015 | 3.111222125 |
| Lrrc32 | -2.982181819 | 2.984625543 |
| Gm18853 | -4.237191002 | 4.353133038 |
| Gm1966 | -8.054247777 | 4.568185492 |
| Gvin1 | -8.063048251 | 4.703439017 |

**Supplementary Table 1. List of significant differentially expressed genes between day four activated CD8 T cells from N2KO vs. N2KO/N1Het.**
